# Forward genetic screening in engineered colorectal cancer organoids identifies novel regulators of metastasis

**DOI:** 10.1101/2023.08.03.551805

**Authors:** Zvi Cramer, Xin Wang, Nicolae Adrian Leu, Keara Monaghan, Joshua H. Rhoades, Yuhua Tian, Joshua Rico, Diego Mendez, Ricardo Petroni, Melissa S. Kim, Rina Matsuda, Maria F. Carrera, Igor E. Brodsky, Ning Li, Christopher J. Lengner, M. Andrés Blanco

## Abstract

Cancer is the second leading cause of death globally, due primarily to metastatic dissemination and colonization of distal sites. Recurrent genetic drivers of metastasis are elusive, suggesting that, unlike the stereotyped mutations promoting primary tumor development, drivers of metastasis may be variable. Here, we interrogate pathways governing metastasis through CRISPR/Cas9-based forward genetic screening in a genetically defined colorectal adenocarcinoma tumor organoid (tumoroid) model using e*x vivo* invasion screens and orthotopic, *in vivo* screens for gain of metastatic potential. We identify *Ctnna1* and *Bcl2l13* as *bona fide* metastasis suppressors. CTNNA1 loss promotes carcinoma cell invasion and migration through an atypical EMT-like mechanism, whereas BCL2L13 loss promotes cell survival after extracellular matrix detachment and non-cell-autonomous macrophage polarization. Ultimately, this study provides a proof-of-principle that high-content forward genetic screening can be performed in tumor-organoid models *in vivo* and identifies novel regulators of colon cancer metastasis.

## Introduction

Colorectal cancer (CRC) represents a major unmet clinical need, accounting for >900,000 deaths worldwide^1^. Despite advances in targeted and immuno-therapies, patients with metastatic disease face poor prognoses. Therefore, it is of great importance to understand the molecular underpinnings of CRC metastasis. Although some drivers of CRC metastasis are known, particularly inactivation of TGF-β signaling^2^, large-scale efforts aimed at identifying recurrent metastasis-specific genetic alterations have been unsuccessful. Thus, novel approaches investigating genes and pathways governing metastasis are needed.

CRISPR-based forward genetic screening enables systematic identification of genes regulating diverse phenotypes and has proven transformative in cancer biology by uncovering critical mediators of cell growth/death, drug resistance, and immunotherapy^4–6^. However, the vast majority of CRISPR screens in cancer research, including CRC, have been conducted in immortalized tumor cell lines *in vitro*. *In vitro* models do not effectively recapitulate the complexity of the metastatic cascade, and findings from cell lines may not translate to primary human cancers. Accordingly, there is a growing need to conduct screens *in vivo* and/or in more physiologically relevant biological models.

Organoid culture of normal, primary epithelia is a powerful technique for *ex vivo* propagation of tissue in a biologically faithful manner. Upon introduction of oncogenic mutations, transformed “tumoroids” can be implanted into syngeneic, immunocompetent mice, where they recapitulate primary tumor growth and the metastatic cascade comparable to human CRC^18^. Thus, organoid/tumoroid models represent a highly attractive setting for CRISPR screening. However, to date few CRISPR screens in organoid/tumoroid models have been conducted, particularly *in vivo*, likely due to the costs and technical challenges associated with these systems.

Here we develop an experimental paradigm for high-content forward-genetic CRISPR screening in CRC tumoroids, assaying invasion and migration *in vitro* and metastasis *in vivo*. We identify novel suppressors of metastasis, including a suppressor of collective cell migration and a regulator of anchorage-independent growth and polarization of tumor-associated macrophages. Thus, we provide a versatile framework for conducting CRISPR screens in organoid cultures that we leverage to identify genes that specifically regulate CRC metastasis.

## Results

### *In vitro* tumoroid screen for regulators of migration and invasion

We set out to apply CRISPR screening to identify novel regulators of CRC metastasis in a biologically faithful tumor organoid system. Accordingly, we generated Cas9-expressing *Apc*, *Trp53*, *Kras*^G12D^ triple mutant mouse organoids (hereafter 3018 *APK*^18^ tumoroids) from normal primary mouse colonic epithelium. These tumoroids can be implanted orthotopically into syngeneic mice, providing a biologically accurate model of colorectal adenocarcinoma with low metastatic proclivity^16^. To facilitate high content screening, we employed methodology^23^ allowing interconversion between 2D and 3D organoid growth (Figure S1A). 2D monolayers proliferated similarly to 3D organoids and could be readily transduced at efficiencies required for screening (Fig S1B and S1C).

We initially set out to screen for genes regulating migration and invasion through the local extracellular matrix (ECM), a critical step in the metastatic cascade, utilizing transwell Matrigel® invasion assays. To optimize screening conditions, we generated clonal *APK* lines with inactivating mutations in the known invasion suppressor *Smad4*^2^ by transient introduction of *Smad4* sgRNAs and Cas9 (hereafter *APKS*). 20-hour assays in 5% Matrigel® resulted in *APKS* cells readily invading through the Matrigel® while *APK* cells did not (Fig S1D). To test our ability to detect clonal gain-of-invasion events, we infected *APK* cultures with lentiviral *Smad4* sgRNAs and identified invasive clones, confirming that this system can detect changes in invasive capacity in response to bulk sgRNA transduction (Fig S1D).

We carried out invasion screens using a curated ‘cancer and apoptosis’ sgRNA library (16,454 sgRNAs targeting 1,562 genes; Table S1)^24^. We transduced APK cells with the library at ∼0.3 MOI and >1,000X coverage and seeded into transwell plates. Cells were collected from the membrane and bottom chamber (invasive), top chamber (non-invasive), and baseline population (used for seeding), and sgRNAs were quantified via next generation sequencing (Fig 1A). The screen was technically successful, with evenly-distributed read counts in the baseline and chamber top populations, and the expected skewing in the bottom chamber consistent with positive selection of phenotypic (invasive) variants (Fig 1B and Table S2). MAGeCK^25^ analysis identified 98 genes significantly enriched in the invasive cells at p ≤ 0.05, 42 of which had ≥ 2-fold median increase in sgRNA counts (Fig 1C, Table S3). Reassuringly, these hits included known migration and invasion suppressors such as *Dcc*^26,27^ and *Ep300*^30^ (Fig S1E and S1F). Pathways enriched in the hit list, but not in a random sampling of the genes in the library, included mesenchymal to epithelial transition (MET) and several morphogenesis pathways, consistent with gene inactivation promoting mesenchymal/invasive phenotypes (Fig 1D and S1G). Thus, this tumoroid screen was technically and biologically successful in identifying regulators of invasion and migration *in vitro*.

**Figure 1.**
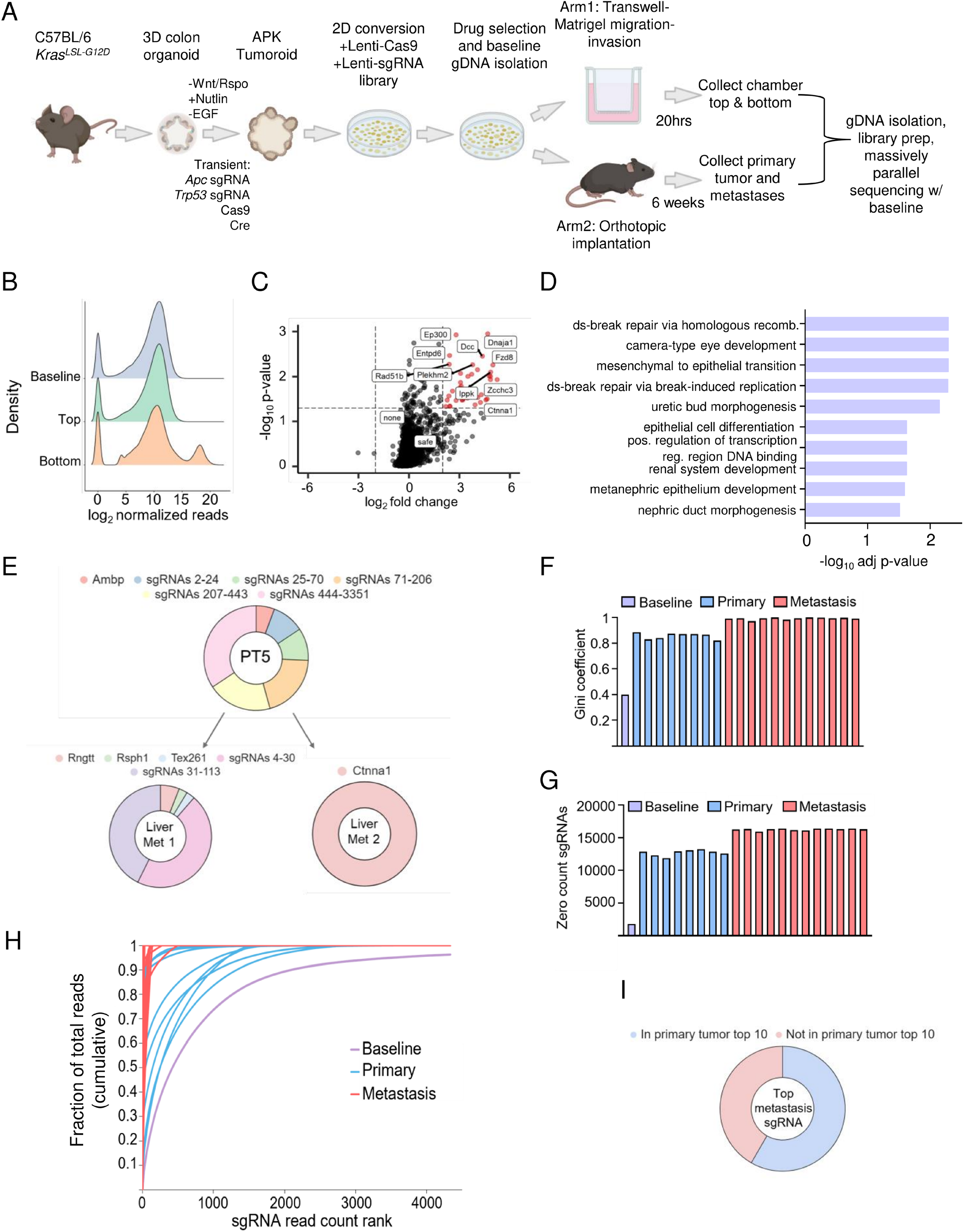
*In vitro* and *in vivo* CRISPR/Cas9 screening of *APK* tumoroids identifies suppressors of invasion and metastasis. **A.** Schematic outline of CRISPR/Cas9 screening platform utilizing monolayer culture. **B.** Density of log_2_ normalized read counts of library sgRNAs in transwell screen. **C.** Volcano plot highlighting top hits (log_2_FC > 2, *p <* 0.05) enriched in transwell membrane relative to top called by MAGeCK as well as negative controls (*sgSafe* and *sgNone*). **D.** GO pathway analysis of enriched hits (*p <* 0.05) in chamber membrane/bottom. **E.** Representative sgRNA makeup of primary tumor and matched metastases. **F,G.** Gini coefficients **(F)** and number of library sgRNAs with 0 reads **(G)** in baseline, primary tumor and metastasis samples. **H.** Cumulative sgRNA read count distribution of baseline, primary tumor, and metastasis samples. **I.** Proportion of metastatic lesions with top sgRNA also found within the top 10 sgRNAs of cognate primary tumors.

### *In vivo* tumoroid screen for regulators of CRC metastasis

As the metastatic cascade cannot be fully modeled *in vitro*, we performed a series of *in vivo* gain-of-metastasis screens (Fig 1A). We delivered the same sgRNA library to APK tumoroids, cultured cells for two weeks, then implanted 1×10^7 cells into the cecum in each of 8 mice, achieving ∼625x library coverage per mouse. Mice were sacrificed 6 weeks post-implantation, a timepoint at which parental cells do not appreciably metastasize. We harvested primary tumors (PTs) and individual, dissected metastatic lesions from the liver, lymph nodes, peritoneal space, and abdominal cavity of each mouse. We quantified sgRNA abundancies from all samples along with the pre-implantation baseline, and first analyzed global characteristics of the PTs (Fig S1H and Tables S4 and S5). As expected given inevitable bottlenecking upon implantation and 6 weeks of *in vivo* growth, library representation was highly skewed in all PTs, with only the baseline sample having a Gini coefficient (a metric of the evenness of sgRNA distribution) < 0.4 (Fig S1I). The degree of skewing varied widely among PTs, with ∼450 sgRNAs accounting for 50% of the total reads in the most diverse tumor, and one sgRNA alone accounting for 53% in the least diverse (Fig S1J). Additionally, 11% of sgRNAs in the baseline were zero counts (no reads detected), while 72%-80% were zero counts in the PTs (Fig S1K). Overall correlation between PTs was low to moderate (Fig S2A), consistent with prevailing dogma that primary tumor progression involves continuous clonal competition leading to marked divergence from the initial sgRNA distribution over time^33^.

We next investigated genes implicated in regulating primary tumor growth in our screen. We focused on sgRNA enrichments in the PTs, as the decreased sgRNA diversity limits the power to detect negatively selected dropouts, but increases the power to identify positively selected tumor suppressor genes (TSGs). We compared each PT to the baseline sample via MAGeCK and found 38 genes were enriched in > 3 PTs at *p* < 0.05 (Table S6). Excitingly, among the strongest were the known tumor suppressor genes *Nf1* and *Nptx2* (Fig S2B and S2C). Altogether, these results support the efficacy of our screening methodology in PTs, both technically and biologically.

We then examined the metastatic lesions. Consistent with expectations, as metastases are thought to be seeded by single of small clusters of cells, we observed dramatically reduced sgRNA diversity in the lesions. After filtering out background sgRNA signal (see methods), metastases had higher Gini coefficients and more zero count sgRNAs than the baseline and PT samples (Figs 1F and 1G). Moreover, metastases exhibited the steepest distribution curves (most skewing), the baseline sample had the most equal sgRNA representation, and PTs largely fell between, with moderate skewing (Fig 1H and Fig S2D).

Our screen provides a unique opportunity to address a longstanding question: whether programs driving metastasis can be disentangled from those driving PT formation^36^. If the phenotypes driving metastasis and PT growth are similar, we would expect sgRNA composition of the PTs to reflect that of its metastatic lesions. However, this was often not the case. In 5 of 12 metastases, the most abundant sgRNA was not among the top 10 PT sgRNAs (Fig 1E and Fig 1I). One mouse was particularly informative, as it had a high-diversity PT that generated 5 metastatic lesions – the most of any mouse. Two of these lesions had one highly dominant sgRNA, with 91% of reads mapping to *Impk* in one and 77% mapping to *Mllt1* in the other – and with neither sgRNA having >1% of the reads in the PT (Fig S2E, Liver Met 1 & 2). The other three lesions were more polyclonal. One did resemble the PT, with the 4 most abundant sgRNAs in the PT all being within the top 5 most abundant in the metastatic lesion (Fig S2E, Liver Met 3), while the other two were dissimilar from the PT, with none of the top 5 sgRNAs in the PT being within the top 5 in the lesion (Fig S2E, Liver Met 4). These data suggest that many, but not all, metastases are generated by sgRNAs conferring a metastasis-specific phenotype that is not selected for in the primary tumor.

### Secondary *in vivo* metastasis screens with focused sub-libraries

As the initial screening library was quite large, we generated a focused sub-library for secondary screens to enable high resolution evaluation of sgRNAs enriched in the metastases compared to PTs. To minimize false negatives, we included all genes with sgRNAs showing any form of enrichment in a metastatic lesion compared to its PT in any mouse (see methods). This liberal criterion yielded 144 genes, to which we added *Smad4* (as a positive control that should promote metastasis) and 10 genes that scored only in the *in vitro* screen. This totaled 465 sgRNAs (3 sgRNAs/gene), to which we added an equal number of safe/non-targeting control sgRNAs to generate a 930 sgRNA library (Table S7).

Secondary screens were performed analogously to the primary screen using an *APK* line derived from a different mouse (3030 *APK*) to avoid clone-specific hits. We obtained and sequenced 7 PTs and 23 metastatic lesions. sgRNA diversity was again decreased in PTs compared to the baseline, and further decreased in the metastatic lesions (Fig 2A and 2B, Fig S3A-E, Fig S4A and B; Tables S8-S10). 13 of 23 lesions were monoclonal/oligoclonal (>70% of reads mapping to one sgRNA), and again the top sgRNAs in the metastases were often not among the top in the PT (Fig S4C and S4D). Critically, in addition to various candidate sgRNAs, sgRNAs targeting the positive control, *Smad4,* were among the most enriched in metastases (Fig S4E and S4F). *Smad4* sgRNAs seeded 3 of 23 metastatic lesions near-monoclonally, and were also moderately enriched in the PTs compared to the baseline (Fig S3F and S3G). This is consistent with reports that *Smad4* can suppress PT growth in addition to metastatic spread^14^, and biologically validates our screening methodology.

**Figure 2.**
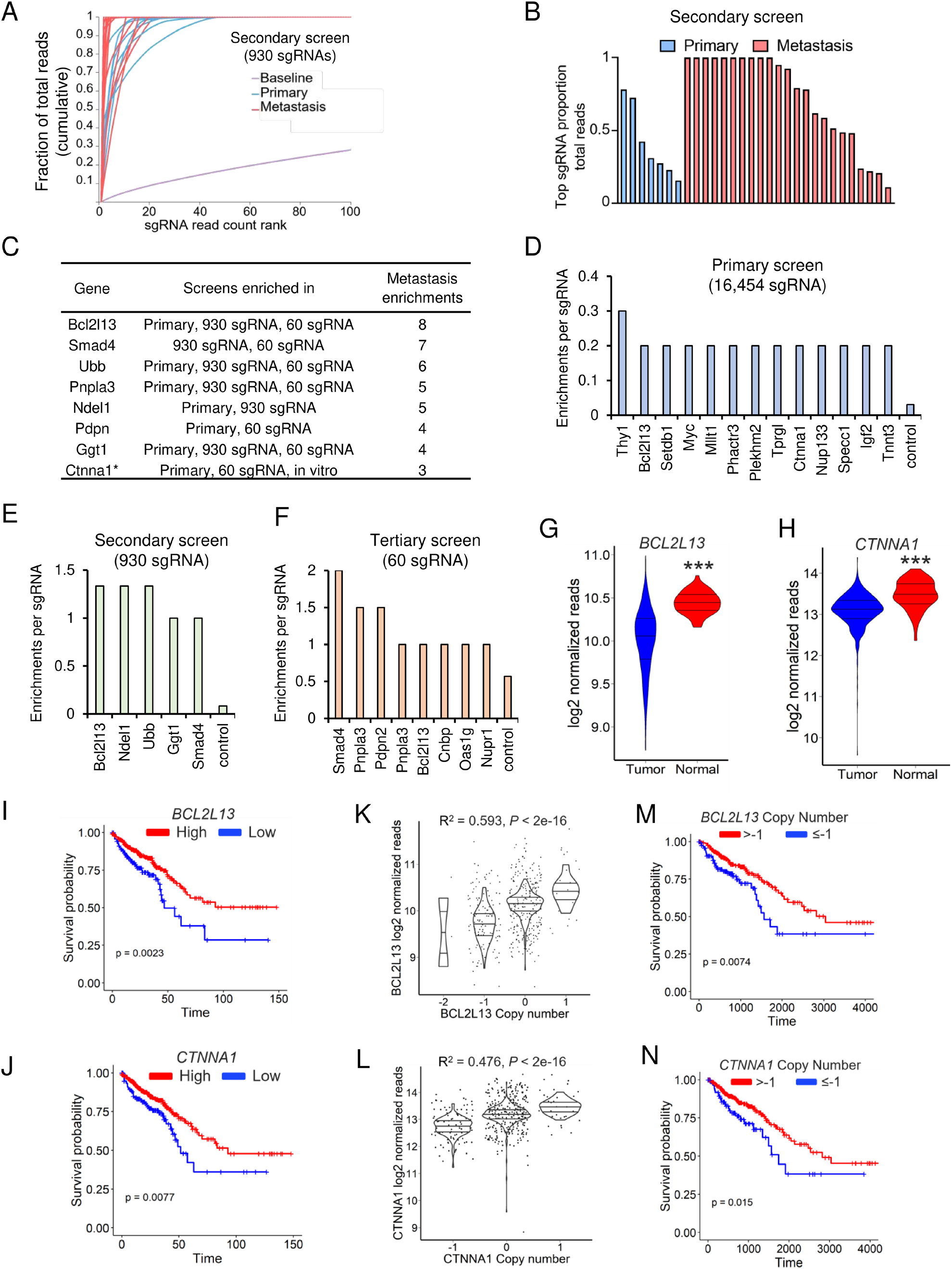
*In vivo* pooled validation of metastasis suppressors, final composite screen hit list and patient survival analyses. **A.** Cumulative sgRNA read count distribution of baseline, primary tumor, and metastasis samples. **B.** The proportion of the total reads mapped to the highest-ranked sgRNA of each primary tumor and metastatic lesion **C.** Table depicting genes with recurrent enrichment in metastases across primary, secondary, and tertiary screens. **D-F.** Number of metastasis enrichments per sgRNA in primary **(D)**, secondary **(E)** and tertiary screens **(F)**. **G,H.** Expression levels of *BCL2L13* **(G)** and *CTNNA1* **(H)** in normal and tumor samples from COAD/READ TCGA datasets. Wilcoxon rank sum test with Holm-Šídák correction was used to assess statistical significance. **I,J.** Overall survival of COAD/READ patients stratified by *BCL2L13* **(I)** and *CTNNA1* **(J)**. Patients were stratified into lowest quartile (low, blue) or highest three quartiles (high, red) and statistical significance was assessed via Mantel-Haenszel tests. **K,L.** Pearson correlation of expression and copy number of *BCL2L13* **(K)** and *CTNNA1* **(L)** in COAD/READ patients. **M,N.** Overall survival of COAD/READ patients stratified by *BCL2L13* **(M)** and *CTNNA1* **(N)** copy number and tested for statistical significance via Mantel-Haenszel test.

We further investigated the relationship between library size and screen performance utilizing a third, highly focused library. We generated a 60 sgRNA library that included 30 sgRNAs targeting selected primary screen hits (and *Smad4*) and 30 safe/non-targeting sgRNAs (Table S11). Analogous screens with this library were technically and biologically sound, with *Smad4* sgRNAs again seeding several near-clonal lesions (Fig S5A-F; Tables S12 and S13). However, the PTs also had markedly decreased diversity. Two of five were essentially clonal, and in the other three the top sgRNA accounted for 40-50% of the reads (Fig S5B). Low diversity in PTs reduces the power of the screen to detect regulators of metastasis, as fewer sgRNAs can compete to seed metastatic lesions. Thus, we conclude that the ∼1,000 sgRNA sub-library size enabled more effective discrimination of regulators of metastasis from regulators of PT growth than the ultra-focused 60 sgRNA library, and was thus a highly effective follow-up to our large-scale primary screen.

We next focused on identifying novel candidate suppressors of metastasis uncovered by our screens for further experimental validation. We analyzed 20 primary tumors and 45 metastatic lesions in total across the primary (16,454 sgRNA), secondary (930 sgRNA), and tertiary (60 sgRNA) screens. Because, to our knowledge, this screening approach is without precedent, hit calling criteria have not been established. The model also presents statistical challenges, as the sample sgRNA abundances do not fit any common distribution [*e.g.*, normal, exponential, or power law (Fig S4G and S4H)], and it is difficult to define replicates, with different mice generating different numbers of lesions. Accordingly, we applied a simple, stringent metric and defined hits as genes with ≥ 3 sgRNA that were strongly enriched in lesions compared to the PT in at least 2 of 3 screens, and were not hits in any PTs (see methods for details). Excitingly, this yielded 8 genes (Fig 2C and Table S14), none of which (outside of the positive control *Smad4*) is an established regulator of metastasis, and only one of which, *Ctnna1,* also scored in the *in vitro* invasion screen. We also observed cases of control sgRNA enrichment, though at markedly reduced frequency compared to gene-targeting sgRNAs after adjusting for number of sgRNAs in the library (Fig 2D-2F). Altogether, these results uncover novel candidate CRC metastasis suppressor genes, and underscore the importance of *in vivo* screening in metastasis studies.

### Confirming and characterizing screen hit phenotypes

We next used CRC patient metadata to identify hits with most potential relevance in human CRC. Among top hits, low expression of *BCL2L13* and *CTNNA1* was the most strongly associated with worse survival of CRC patients, and both genes are upregulated in normal tissue compared to tumors (Figs 2G-J). Remarkably, we also identified partial copy number loss in both *BCL2L13* and *CTNNA1* (31.9% and 23.1%, respectively) in COAD/READ patients (Fig S6A and S6B), and copy number loss was significantly associated with decreased gene expression and worse patient survival (Figs 2K-N). *CTNNA1* and *BCL2L13* were also identified as copy number dosage-sensitive genes in CRC^41^, and multiple studies identified recurrent deletions in the chromosomal region 22q11 encompassing *BCL2L13*^19,42–44^, with one study directly associating 22q11 loss with liver metastasis incidence^44^. Finally, both genes reside on chromosome arms lost as late events in CRC^45^. Thus, alterations of BCL2L13 and CTNNA1 are frequent in CRC patients and strongly associated with worse survival.

We selected *Bcl2l13* and *Ctnna1* for individual validation. We delivered sgRNAs targeting these genes (Fig S6C and D) or control genomic regions (*sgSafe*) to 3018 *APK* tumoroids and implanted bulk populations into syngeneic mouse ceca. After 5-6 weeks, PTs and metastases were harvested for analysis. Knockout of neither gene significantly affected PT weight, proliferation (Ki67+), or apoptosis (cleaved-caspase-3+) of carcinomas (Fig 3A and 3B, Fig S6E-S6H). However, loss of *Bcl2l13* or *Ctnna1* both markedly increased liver macrometastasis incidence (Fig 3C-3E), with *Ctnna1* KO lesions appearing poorly differentiated (Fig S6E). *In vitro,* inactivation of *Bcl2l13* or *Ctnna1* did not affect APK tumoroid cell death, though, interestingly, *Ctnna1* loss decreased the proportion of active S-phase cells (Fig S7A-S7D). Excitingly, these findings are in close agreement with screening results, and confirm *Ctnna1* and *Bcl2l13* as novel suppressors of *APK* metastasis that do not regulate proliferation or cell death in the PT.

**Figure 3.**
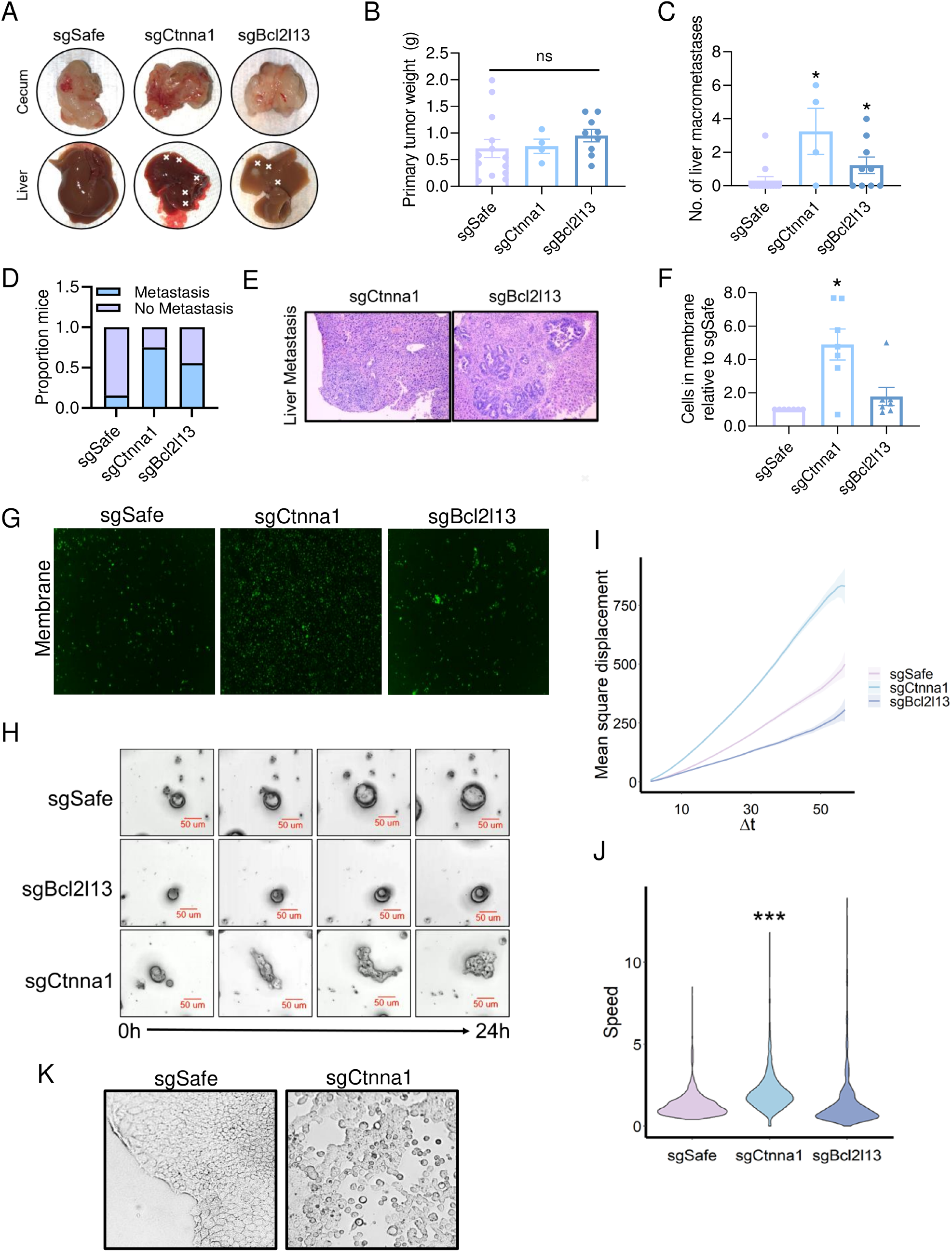
Candidate metastasis suppressor genes regulate metastasis but not primary tumor growth. **A.** Representative images of primary tumor and liver metastases formed after implantation of *APK* tumoroids transduced with the indicated sgRNAs into the cecum of syngeneic mice. **B-D.** Size of primary tumors **(B)**, number **(C)** and frequency **(D)** of liver metastases formed by orthotopically implanted *APK* organoids transduced with the indicated sgRNAs. Two sgRNAs targeting each gene were combined for each experimental group (*n ≥* 4 each group) and statistical significance was evaluated via Mann-Whitney test with Holm-Šídák correction. Error bars represent mean ± SEM. **E.** Representative hematoxylin and eosin-stained sections derived from liver metastases formed by orthotopically implanted *APK* organoids transduced with the indicated sgRNAs. Scale bar= 250 µm. **F,G.** Quantification **(F)** and representative images **(G)** of Hoechst-stained *APK* tumoroids transduced with *sgSafe*, *sgCtnna1-1* or *sgBcl2l13-1* in transwell membranes 20h post-seeding. Fiji was used to quantify signal in membranes. Statistical significance was assessed by student’s paired T-test with Holm-Šídák correction. (*n =* 7). Error bars represent mean ± SEM. **H.** Representative stills from live imaging of *APK* tumoroids transduced with *sgSafe*, *sgCtnna1-1* or *sgBcl2l13-1* and seeded on top of diluted Matrigel. Scale bar = 50µm. **I,J.** Mean square displacement **(I)** and speed **(J)** of individual tumoroids in arbitrary units during 24h of live imaging. Movement tracks and quantification determined by Fiji and celltrackR, respectively. Statistical significance was assessed by ANOVA with Tukey’s post hoc test. (*n ≥* 176). **K.** Brightfield images of *APK* monolayers transduced with *sgSafe* or *sgCtnna1-1*. Scale bar = 500µm. For all panels * *p* < 0.05, *** *p* < 0.001.

We next investigated the mechanisms by which *CTNNA1* and *BCL2L13* suppress metastasis. CTNNA1 is a tension-dependent scaffold between adherens junctions and the actin cytoskeleton that has been implicated in cell-cell adhesion and regulation of signaling pathways^46^. BCL2L13 is a BCL2-like outer mitochondrial membrane protein that promotes PINK1/PARKIN-independent mitophagy and has both pro- and anti-apoptotic functions^47,48^. We first tested whether these proteins suppress tumoroid invasiveness. Consistent with *in vitro* screen data, *Ctnna1* KO induced a dramatic, nearly 5-fold increase in invasiveness in transwell assays, while *Bcl2l13* KO had no effect (Fig 3F and 3G). Remarkably, live imaging of tumoroids revealed that *Ctnna1* KO also induced extensive and rapid collective cell migration with leading edge protrusions, while *Bcl2l13* KO and control tumoroids remained relatively immobile (Fig 3H-J, Videos 1-3). Given these phenotypes and the reduced cell-cell adhesion in *Ctnna1* KO monolayers (Fig 3K), we asked whether CTNNA1 loss promotes EMT (epithelial-to-mesenchymal transition),which is often associated with gain of invasiveness, metastasis, and cancer stem cell (CSC) activity^49^. In line with this, *Ctnna1* KO increased tumoroid formation ability, a proxy for tumor initiating or CSC activity (Fig S7G and S7H). To our surprise, *Ctnna1* KO cells maintained surface E-cadherin expression *in vivo*, and did not have changes in the canonical EMT markers *Cdh1*, *Cdh2*, *Snail1*, *Twist*, *Vim*, *Zeb1*, and *Zeb2 in vitro* (Fig S7E and S7F). Altogether, these data suggest that loss of CTNNA1 promotes metastasis by increasing both the migratory and invasive ability of APK tumoroids in the absence of canonical EMT and in association with elevated CSC activity.

BCL2L13 is a known apoptosis regulator, and its loss did not confer a migrative/invasive phenotype. Thus, we tested whether it could affect apoptosis or anoikis, a specialized form of apoptotic cell death induced by detachment from the ECM that metastatic cells must suppress to survive in circulation. Intriguingly, while *Bcl2l13* KO did not affect apoptosis in native, adherent culturing conditions, it dramatically and consistently suppressed cell death upon loss of adherence (Fig 4A-4C, Fig S8A and S8B). Interestingly, this survival phenotype appeared independent of apoptosis, as caspase inhibition did not phenocopy BCL2L13 loss (Fig S8C). As BCL2L13 regulates mitophagy, a process that can induce non-apoptotic anoikis^50,51^, we tested whether its loss affected aspects of mitophagy. Indeed, *Bcl2l13* KO increased the mitochondrial DNA:nuclear DNA (mtDNA:nDNA) ratio and the level of the outer mitochondrial membrane protein TOM20 (Fig 4D-4G), both of which are indicators of mitophagy inhibition. In the context of metastasis, these findings suggest that BCL2L13 loss may prevent cell death after delamination from the primary tumor, thereby allowing cells to survive in circulation until ECM re-engagement.

**Figure 4.**
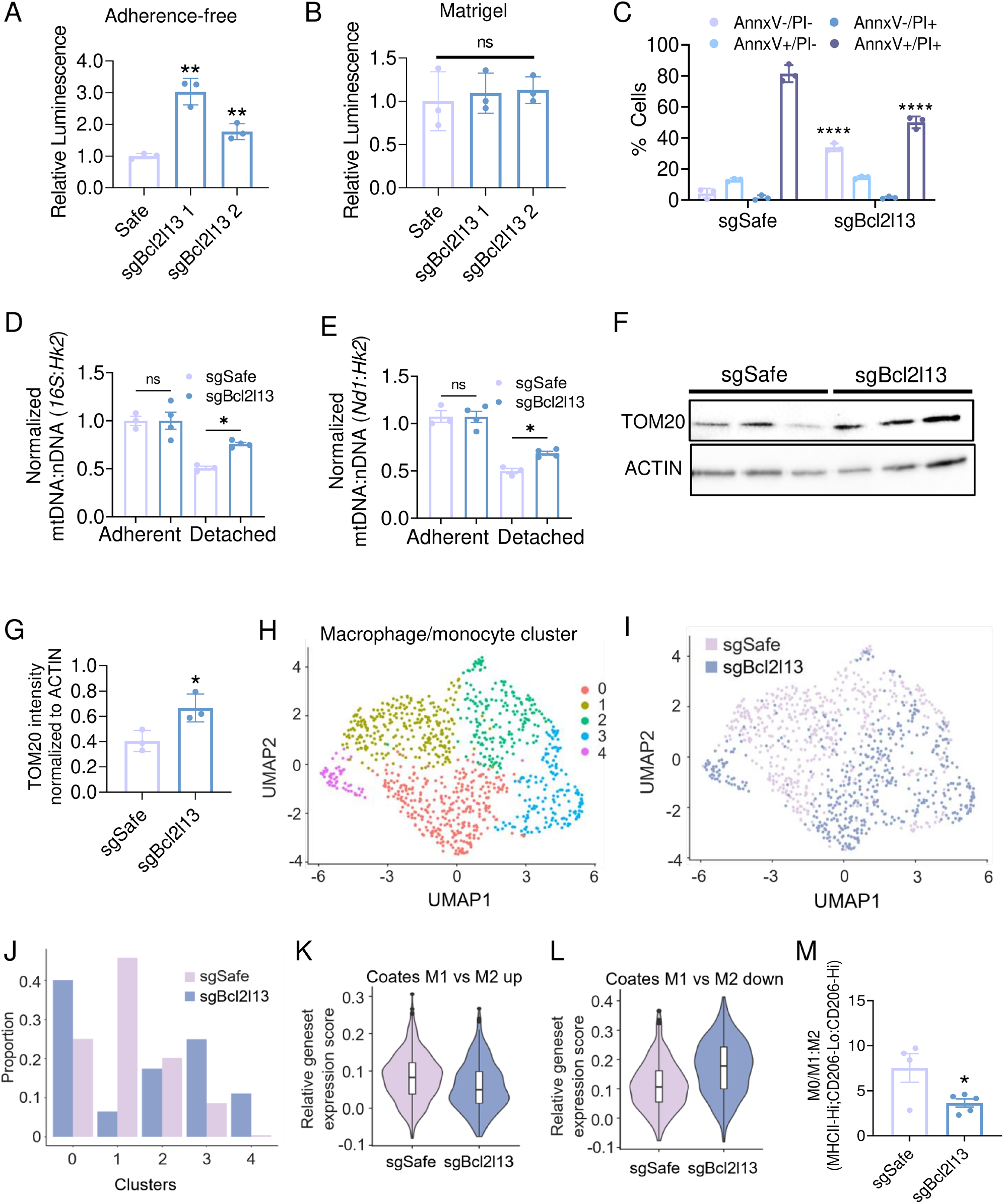
BCL2L13 regulates survival after extracellular matrix detachment and macrophage polarization. **A, B** CellTiter-Glo assay luminescence normalized readings from *APK* tumoroids transduced with the indicated sgRNAs after 72h growth in detached **(A)** or adherent **(B)** conditions. (*n*= 3). Error bars represent mean ± SD. **C.** Quantification of Annexin V and propidium iodide (PI) staining of *APK* tumoroids transduced with *sgSafe* or *sgBcl2l13-1* after 24h detachment. (n = 3). Error bars represent mean ± SD. **D,E.** Ratio of mtDNA-encoded genes *16S* **(D)** and *Nd1* **(E)** in *APK* tumoroids transduced with *sgSafe* and *sgBcl2l13-1* and grown in adherent or detached conditions for 24h. (*n ≥* 3). Error bars represent mean ± SEM. **F,G.** Western blot **(F)** and quantification **(G)** of the mitochondrial protein TOM20 in *APK* tumoroids transduced with *sgSafe* and *sgBcl2l13-1* and grown for 24h in detached conditions. β-ACTIN was used as a loading control. (*n ≥* 3). **H,I.** Uniform Manifold Approximation and Projection (UMAP) for macrophage/monocyte populations profiled from *sgSafe* and *sgBcl2l13-1 APK* primary tumors colored by cluster **(H)** and genotype **(I)**. **J.** Proportion of monocytes/macrophages in each cluster shown in Fig 6H. **K,L.** Relative gene set expression score of M1-like genes **(K)** and M2-like genes **(L)** calculated using Seurat. **M.** M0/M1:M2 ratios in individual *sgSafe* and *sgBcl2l13-1/sgBcl2l13-2 APK* primary tumors. M0/M1 and M2 macrophages were defined as MHCII-Hi;CD206-Lo and CD206-Hi macrophages respectively (*n ≥* 4). *sgBcl2l13-1* and *sgBcl2l13-2* were combined. Error bars represent mean ± SEM. For panels H-L two biological replicates for each genotype are merged. For panels (A), (B), (G), and (M), statistical significance was assessed via student’s t-test, with Holm-Šídák corrections applied in (A) and (B). For panels (C-E), statistical significance assessed via two-way ANOVA with Holm-Šídák multiple comparisons correction. For all panels, * *p* < 0.05, ** *p* < 0.01, **** *p* < 0.0001.

To further investigate the mechanisms by which BCL2L13 suppresses metastasis, we performed single cell RNA-seq (scRNA-seq) on *sgSafe* and *sgBcl2l13 APK* PTs. UMAPs (Uniform Manifold Approximation and Projection) identified clear clusters of carcinoma cells, normal colonic epithelium, B and T cells, fibroblasts, and several myeloid cell types (Fig S8D and S8E). GSEA indicated that *sgBcl2l13* carcinoma cells downregulated the interferon alpha and gamma (IFN-α/ IFN-γ) response gene signatures and upregulated the hypoxia response (Fig S8F and S8G). In line with our experimental results, *Bcl2l13* KO did not affect proliferation-related gene sets (Figs S8G).

We next investigated RNA-seq data in stromal cells. While the abundances of most stromal cell types were similar in *sgSafe* and sg*Bcl2l13* PTs, we observed dramatic differences in the distribution of macrophage/monocyte cell subtypes (Figs 4H-J, Fig S9A). Macrophages can be broadly classified as M1-like or M2-like according to their reactive functionality, or polarization. M1-like are associated with the inflammatory response and inhibition of tumor progression, while M2-like are considered tumor and metastasis-promoting through immune-suppression, angiogenesis, and microenvironmental remodeling^5253^. Intriguingly, M2-like gene signatures and classic M2 markers were strongly enriched in macrophages from *sgBcl2l13* PTs, while M1-like signatures, associated pathways, and markers were predominant in *sgSafe* PT macrophages (Figs 4K and 4L, Figs S9B-F). Notably, the secreted factors *Vegfa* and *Ccn2* (CTGF), both drivers of M2 polarization, were also elevated in *sgBcl2l13* carcinoma cells^54,55^ (Fig S9G). To verify our scRNA-seq results, we performed flow cytometry on *sgSafe* and *sgBcl2l13* PTs (Fig S10A). Indeed, we found a higher ratio of M2-like macrophages in *sgBcl2l13* PTs without changes in total macrophage infiltration (Fig 4M and Figs S10A-10C). Taken together, these results suggest that M2 macrophage polarization may also contribute to metastatic suppression by BCL2L13.

Altogether, this study provides a framework for utilizing high-content forward genetic screening using tumor organoids in an *in vivo* model for carcinoma metastasis and identifies novel metastasis suppressors in colorectal adenocarcinoma.

## Discussion

There is a great need to identify regulators of metastasis, and unbiased genetic screening provides an opportunity to do so. However, efforts to date have generally screened cancer cell lines in immunodeficient mice and quantified metastasis sgRNA abundance in whole target organ lysates (*i.e.*, not individual metastatic lesions). A small number of organoid-based *in vivo* CRC screens have also been reported^20–22^, but these have focused only on primary tumor formation and utilized small sgRNA libraries. Here, we generated and leveraged methodology for performing high content *in vitro* and *in vivo* CRISPR screens in genetically-defined CRC organoids/tumoroids.

Using our screening pipeline, we identified and experimentally confirmed CTNNA1 and BCL2L13 as novel metastasis suppressor genes that function through distinct mechanisms. CTNNA1 restrains cell self-renewal, migration and invasion programs, and its loss induces a mesenchymal-like morphology in the absence of a classical EMT. In contrast, BCL2L13 loss helps prevent non-apoptotic cell death upon ECM detachment. In addition to these cell-autonomous phenotypes, we also found increased M2-like macrophage polarization in *sgBcl2l13* tumors. M2-like macrophages are associated with almost every step of the metastatic cascade^53^, suggesting changes in macrophage identity could further underlie the suppression of metastasis by BCL2L13. As clinical data also implicate CTNNA1 and BCL2L13 in CRC metastasis, further study of their mechanisms of action may uncover novel opportunities for therapeutic intervention in patients with metastatic CRC.

Our screening approach has broader implications for understanding the metastatic process. An outstanding question is whether there exist genes that specifically regulate metastasis independently from primary tumor growth. CRC genome sequencing studies have found relatively few mutations linked to metastasis but not primary tumor formation, suggesting that gain of metastatic ability is either a low probability stochastic event or is a more general feature of aggressive primary tumor cells. However, a number of genes have been shown to regulate metastasis with little or no effect on the primary tumor in experimental mouse models^66^. Our results provide support for both possibilities. Loss of CTNNA1 and BCL2L13 markedly increased metastatic ability without having any significant effect on primary tumor growth. However, the observed cases of non-targeting sgRNA enrichments in metastatic lesions indicates that metastatic ability can also be acquired via presumed non-genetic clonal evolution^31^.

Our approach has certain limitations. Extensive cell death known to occur^31,32^ upon orthotopic transplantation dramatically reduces sgRNA diversity at an early point. There is also a small number of positive events (ie. metastatic lesion) relative to library size, suggesting almost certainly that our screens had a large number of false negatives. In future studies this can be at least partially mitigated by increasing the number of mice used and/or using cells with a slightly increased basal metastatic ability.

Finally, the screening methodology developed here is flexible and scalable. Increasing screen scale may enable the identification of organ-specific metastasis regulators, and inducible Cas9 models could uncover regulators of specific stages of metastasis. Analogous screens performed with CRISPR activation systems could identify drivers of metastasis, rather than suppressors. This approach in particular may generate results of high therapeutic interest.

## Acknowledgments

The authors thank the following core facilities: Penn Flow Core, Center for Molecular Studies in Digestive and Liver Diseases (NIH P30-DK050306) Molecular Pathology and Imaging Core, Penn Vet Imaging Core (Gordon Ruthel). This work was supported by the following: NIH F31CA250267-02 (ZC), NIH NRSA F31AI160741-01 (RM), a Calico Labs sponsored research agreement (CL and MAB), and NIH/NCI R01 CA279317-01 (MAB).

## Declaration of Interests

The authors declare no competing interests.

## Author Contributions

Conceptualization: M.A.B., C.J.L., Z.C., X.W., I.E.B., and N.L.

Experimentation: Z.C., X.W., N.A.L., K.M., Y.T., J.R., D.M., R.P., N.L., M.S.K., and R.M.

Computational and statistical analyses: J.H.R., M.A.B., Z.C., X.W., and M.F.C.

Writing: M.A.B., C.J.L., Z.C., X.W., and M.F.C.

## Star Methods

CONTACT FOR REAGENT AND RESOURCE SHARING

Request for more information about this manuscript and any reagents or codes used therein will be fulfilled upon request to the Lead Contact, M. Andrés Blanco

## Organoid and monolayer derivation

Murine colon organoids were cultured as previously reported (Sato et al., 2009). Briefly, the colons of LSL*-Kras*^G12D^ mice were removed, scraped lightly, washed with cold D-PBS until clean and then digested with 5 mM EDTA (Invitrogen #15575-020) for 20 min at 4°C. Crypts were released by scraping and then filtered through 70µm cell strainer to remove tissue fragments. For organoid culture, crypts were resuspended in 3:1 Matrigel:PBS, seeded into 24-well plates at a density of ∼500 crypts in 50 µl total volume per well and incubated with complete medium. Complete medium was composed of Advanced DMEM/F-12 (Thermo Fisher Scientific #12634010) containing 1X B27 (Gibco #17504-044), 1X N2 (Invitrogen #17502-048), 1X GlutaMAX (Thermo Fisher Scientific #35050061), 10mM HEPES (Thermo Fisher Scientific #15630-080), 1mM N-acetyl-cysteine (Sigma-Aldrich #A9165), supplemented with 1x conditioned media (derived from L-WRN cells). Normal organoid media was also supplemented with 50 ng/ml mEGF (Invitrogen #PMG8041). Monolayer cultures were derived as previously described (Wang 2017). Collagen I (Corning #354236) was first diluted in collagen neutralization buffer (final concentration: 1 mg/mL collagen I, 1X PBS, 22 mmol/L HEPES, 55 mmol/L NaHCO_3_, 0.006N NaOH). 1mL of diluted collagen was added to 6-well plates and allowed to polymerize for 1 hour at 37°C. ∼1500 crypts were added to each well in 1mL complete medium supplemented with 500 nmol/L A83-01 (Tocris #2939). Both organoids and monolayers were treated with 10µM Y-27632 (Selleck Chemicals #S1049) for 2 days after seeding.

## Passaging and maintenance of organoid/tumoroids and monolayer cultures

For organoids/tumoroids, cultures were allowed to grow for 5-7 days at 5% O_2_ and media was changed every other day. Organoids were collected in Dispase (STEMCELL Technologies #07923) and incubated for 45 min at 37°C to release organoids from Matrigel. Organoids were then washed with PBS, pelleted and resuspended in Accutase (Sigma-Aldrich #A6964). During maintenance passaging to 3D or 2D, organoids were incubated in Accutase for 5 min at 37°C. 1mL 5% BSA PBS was then added to the organoids which were subsequently pipetted into 1-10 cell clusters and washed further with 10mL PBS before pelleting by centrifugation. During single-cell passaging, organoids were incubated in Accutase for 7-8 min at 37°C and vigorously pipetted after addition of 5% BSA PBS. Cells were then washed further with 10mL PBS and pelleted by centrifugation for counting. For both maintenance and single-cell seeding, cells were resuspended in 3:1 Matrigel:PBS and seeded as domes on culture plates. Seeded matrigel domes were allowed to polymerize at 37°C for 10-15 min and then complete medium supplemented with 10µM Y-27632 was added.

Monolayers were passaged as previously described^23^, with some modification. Monolayer cultures were allowed to grow until 70-90% confluence with media changes every other day. Media was removed and collagen hydrogels were collected in a 15mL conical tube in 1mL 500U/mL collagenase IV (Gibco #17104019) per 1mL collagen hydrogel, incubated at 37°C for 10 minutes and subsequently washed with 9mL PBS and pelleted by centrifugation. In the case of larger volumes of collagenase (>4mL), 10X collagen was diluted to 1X directly into the collected gel/residual buffer. For maintenance of monolayer culture, pelleted monolayers were resuspended in 1mL of 0.5mM EDTA-PBS, pipetted vigorously until monolayers were broken into 5-10 cell clusters and washed with 9mL of PBS before pelleting by centrifugation and seeding on top of collagen hydrogels in complete medium supplemented with 500 nmol/L A83-01 and 10µM Y-27632. For conversion into organoid/tumoroid culture, pelleted monolayers were incubated in 500µL Accutase for 5 min at 37°C. 1mL 5% BSA PBS was then added to the organoids which were subsequently pipetted into 1-10 cell clusters and washed further with 10mL PBS before pelleting by centrifugation. Cells were then seeded in 3:1 Matrigel:PBS, allowed to polymerize, and then complete medium supplemented with 10µM Y-27632 was added.

## Genome engineering of mouse organoids

CRISPR/Cas9 editing of organoids was performed in two-dimensional monolayers as previously described^18^. Briefly, the KrasG12D mutation was activated by Cre-mediated deletion of the stop cassette at the Kras-LSL-G12D targeted allele by transfection of a plasmid expressing Cre recombinase along with a pPGK-Puro plasmid (a gift from Rudolf Jaenisch, Addgene#11349) using Lipofectamine 2000 (Thermo Fisher Scientific #11668019) followed with 10 μg/ml puromycin selection for 3 days. The *Apc*, *Trp53* and *Smad4* mutations were introduced by CRISPR-Cas9 editing. sgRNAs targeting *Apc*, *Trp53* and *Smad4* were cloned into pX330 (a gift from Feng Zhang, Addgene #42230). *Apc* and *Trp53* sgRNAs were first introduced into organoids and after one week, *Apc* mutants were selected by withdrawal of L-WRN and addition of recombinant 100ng/mL Noggin (Peprotech 230-38) and *Trp53* mutants were selected for by adding 10 µm Nutlin-3 (Cayman Chemical Company #10004372-1). Resulting *APK* lines were subcloned and then sgRNAs targeting *Smad4* were introduced. *Smad4* mutants were then selected for by withdrawal of L-WRN/Noggin. Genotypes of subcloned tumoroid lines were verified by Sanger sequencing. Subclones derived from mouse 3018^18^ were used in all experiments except pooled validation studies (mouse 3030).

## Mouse models

All animals were maintained on a C57BL/6J background. C57BL/6J (JAX strain #000664) and LSL*-Kras^G12D^* (JAX strain #008179), mice were obtained from The Jackson Laboratory. The Institutional Animal Care and Use Committee of the University of Pennsylvania (Animal Welfare Assurance Reference Number A3079-01, approved protocol no. 803415 granted to Dr. Lengner) approved all procedures involving mice and followed the guidelines set by the Guide for the Care and Use of Laboratory Animals of the National Research Council of the National Institutes of Health.

## Generation of vectors and pooled sgRNA library cloning

The pX330-U6-Chimeric_BB-CBh-hSpCas9 (a gift from Feng Zhang, Addgene #42230), LentiCas9-Blast (a gift from Feng Zhang, Addgene #52962) and LentiGuide-puro (a gift from Feng Zhang, Addgene #52963), which express sgRNA and/or Cas9 nuclease, were used to generate gene-specific sgRNA vectors. For individual sgRNA cloning, pX330 was digested with BbsI (NEB #R3539S) and ligated with annealed sgRNAs targeting *Apc*, *Trp53*, or *Smad4*. LentiGuide-puro was digested with BsmBI (NEB #R0739S) and ligated with annealed sgRNA oligos targeting *Ctnna1* and *Bcl2l13*. The sgRNA target sequences used included: *sgApc*: GTAATGCATGTGGAACTTTG;*sgTrp53*:TCCGAGTGTCAGGAGCTCC;*sgSmad4*: GATGTGTCATAGACAAAGGT;*sgBcl2l13-1*:GATATGCCTCATATGTCAC;*sgBcl2l13-2*:GGTGTTGTAGCAAAACTAG;*sgCtnna1-1*:GATGGTATATTGAAACTG;sg*Ctnna1*-2- :GATTTGATGAAGAGCGCTGC

sgRNA pool design and amplification were performed according to a previous protocol^67^. Briefly, for validation libraries, custom pools were designed using Broad GPP sgRNA design tool and synthesized by Azenta Life Sciences. Pools were then cloned into Bbs-digested LentiGuide-puro backbone using Gibson Assembly (NEB E2611S). Both validation pools and half A of Apoptosis and cancer library (a gift from Michael Bassik, Addgene #1000000121) were amplified using Phusion High-Fidelity polymerase (NEB E0553S). Amplified products were then transformed into electrocompetent cells (Lucigen).

## Lentiviral preparation and infection

Viral vectors were maxiprepped using ZymoPURE II™ Plasmid Maxiprep (Zymo Research D4202) and then transfected with PEI (Polysciences 49553-93-7) into HEK293T cells with packaging plasmids psPAX2 (Addgene #12260) and pMD2G (Addgene #12259). The viral supernatant was collected after 48 hours and 72 hours and passed through a 0.45µm filter. Lenti-X Concentrator (Clontech # 631231) or ultracentrifugation was used to concentrate virus 50-100X. Monolayers on collagen hydrogels were infected with lentivirus plus 1 μg/ml polybrene (Sigma-Aldrich #TR-1003-G) for 6h. Three days after the infection, monolayers were selected with the antibiotics Blasticidin (20 µg/ml) and/or Puromycin (10 µg/ml).

## Transwell assay

Matrigel (Corning #356231) was thawed at 4 °C overnight and diluted accordingly in Matrigel dilution buffer (0.01M Tris pH 8.0, 0.7% NaCl). Transwell plates with a pore size of 8μm (Corning #3428) were coated with 1:20 and 1:30 diluted Matrigel in the upper and lower chambers respectively. The plates were then incubated at 37°C for 2h, after which Matrigel solution was removed. Tumoroids or monolayers were digested to single cells/small cell clusters with Accutase and resuspended in basal medium (Advanced DMEM/F12, 10 mM HEPES, 1× Penicillin-Streptomycin, 2 mM GlutaMAX, 1× N2 supplement, 1× B27 supplement, 1 mM N-acetyl-cysteine) supplemented with 10µM Y-27632, whereas complete media (basal medium supplemented with L-WRN conditioned media and 10µM Y-27632) was added to the bottom chamber. ∼8*10^6 or 0.1-0.2*10^6 cells were seeded into the upper chambers of 6- and 96-well plates respectively and incubated for 20h before collection/analysis.

For CRISPR screening in 6-well plates, the top of the membrane was thoroughly washed and scraped with a pipette tip and then all cells in the top chamber were collected. The transwell membrane and bottom were then incubated at 37°C in Dispase (STEMCELL Technologies #07923) for 30 minutes and was thoroughly washed with 0.5% BSA PBS to dislodge cells. Both top and bottom cells were pelleted and frozen at −80°C for future library preparation. For single-gene validation in 96-well plates, cells on top of the membrane were first removed by washing and scraping, then cells remaining in membrane were fixed with 4% PFA, washed, and stained with Hoescht 33341 (ThermoFisher #H3570). Membranes were then imaged using fluorescent microscope on a Leica DM500 Widefield microscope. Quantification of cell number in the membrane was quantified using Analyze Particles function in Fiji.

## Cecum Implantation

Tumoroids were collected and dissociated into clusters of 3–5 cells. For each mouse, approximately 10 million cells were suspended in 90µL implantation medium (Advanced DMEM supplemented with B27, N2, and 10µM Y-27632) with 10µL Matrigel. 8-week-old male C57BL/6J (JAX strain #000664) were anesthetized with Ketamine/Xylazine and the cecum was exteriorized by performing a midline abdominal incision. Next, under a binocular lens, the cell suspension was injected into the cecal sub-serosa with an insulin needle. Afterwards, the area around the injection site was flushed with PBS, the cecum was then returned to the abdominal cavity which was thereafter closed. The mice were closely monitored for signs of distress until the endpoint of the experiment.

## Screening library preparation and sequencing

For library preparation, genomic DNA (gDNA) from the metastatic tumors from in vivo screening and the cells in the lower chamber from in vitro transwell assay was extract with QIAamp DNA Micro Kit (Qiagen #51306), while gDNA from the primary tumor and the cells in the upper chamber was extracted with DNeasy Blood & Tissue Kit (Qiagen #69504). For both main and validation screens, we designed a two-step PCR protocol for efficient amplification of integrated sgRNAs. For the main screen with Cancer and Apoptosis library, we utilized step 1 PCR primers containing pMCB320 backbone homology sequence, staggers and partial Illumina adaptor sequence (Supplementary File 1) and the step 2 PCR primers contained Illumina sequencing adaptors and barcode indices (Supplementary File 1). Final PCR products were ran on an agarose gel and purified for sequencing using Purelink gel extraction kit (Thermo Scientific K210025). For the validation screening, step 1 PCR primers containing a LentiGuide-puro homology sequence and step 2 primers containing staggers and Illumina adaptors were designed. Final PCR products were then purified by AMPure XP Reagent (BECKMAN #A63880) or gel extraction (Thermo Scientific K210025). Takara ExTaq Polymerase (Takara Bio RR001A) was used for all PCR reactions. After purification, the sequencing libraries were pooled and sequenced using Illumina HiSeq 4000, capturing reads >1000X the number of sgRNAs of the screening library used.

## CRISPR Screen Analyses

Normalized read count files for all screens were generated using the MAGeCK count command with default settings. For *in vitro* transwell screen analyses, enrichment of sgRNAs and genes in the chamber bottom sample compared to the chamber top sample was analyzed with the MAGeCK test command with default settings. Genes that were enriched in the bottom sample compared to the top sample at p < 0.05 and with median sgRNA log_2_ fold change ≥ 2 in the bottom compared to the top sample were considered hits. For *in vivo* screens, enrichment of genes in each primary tumor sample compared to the baseline sample was analyzed with the MAGeCK test command with default settings. Genes that were enriched in a given primary tumor compared the baseline sample at p < 0.05 were considered hits in that tumor. For analyses of enrichment in metastatic lesions, sgRNAs were first filtered out if they had ≤ 1 RPKM in all samples. In the primary screen, all sgRNAs that comprised the bottom 10% of a given metastatic lesion sample were also filtered out, and lesions were discarded if the top 10 most abundant sgRNAs comprised < 20% of the total reads in the given sample. In the secondary and tertiary screens, only the < 1 RPKM sgRNA filter was used in the metastatic lesion samples. In the primary screen, an sgRNA was considered enriched in a given metastatic lesion if it had a fold change of ≥ 100 in the lesion compared to its matched primary tumor and also had a read count of ≥ 25,000 RPKM in the lesion. For sgRNAs that were not detected in the primary tumor, the fold change criterion was removed. Exceptions to these criteria were sgRNA “jackpots,” which were sgRNAs comprising ≥ 70% of the total reads of the metastatic lesion. Jackpot sgRNAs only had to have a fold change of ≥ 3 in the lesion compared to the primary tumor to be enriched. For the secondary and tertiary screens, the same criteria for enrichment in a metastatic lesion compared to its matched primary tumor were used, except that sgRNAs only had to have a fold change of ≥ 25 in the lesion compared to the primary tumor (in addition to having ≥ 25,000 RPKM in the lesion). Considering all screens together, gene-level hits were defined as genes with at least three sgRNAs, each from a different metastatic lesion, enriched compared to its matched primary tumor in at least two of the three screens.

## TCGA survival and copy number variation analysis

For the analysis of the association of *CTNNA1* and *BCL2L13* with human colorectal cancer patient survival, TCGA COAD/READ expression and survival data was obtained from Cbioportal (Colorectal Adenocarcinoma (TCGA, PanCancer Atlas)). Low expression patients for each gene were defined as those in the bottom quartile of gene expression, whereas high expression patients were defined as those in the top three quartiles. For tumor vs normal and copy number variation analysis, gistic-thresholded copy number values and gene expression data was acquired from UCSC Xena^68^ TCGA Colon and Rectal Cancer (COAD/READ) dataset. For all survival analyses, R packages *Survival* and *Survmine* were used. Results were visualized using ggplot2.

## Clonal tumoroid formation assays

Tumoroids were digested into single cells and filtered through a 40µm filter. Viable cell numbers were determined using a hemocytometer and equal numbers of single viable cells were seeded in 24 well plates with Matrigel at a density of 750 cells/well in complete medium supplemented with 10µM Y-27632. Brightfield images of each well were captured 4 days post-seeding with a Leica DM500 Widefield microscope and manually counted using Fiji.

## Immunofluorescence

After dissection and washing with PBS, tumors were fixed in 4% paraformaldehyde (Electron Microscopy Sciences #15710), paraffin-embedded and sectioned. Haematoxylin & eosin staining was conducted by Morphology Core of the UPenn NIDDK P30 Center for Molecular Studies in Digestive and Liver Diseases. For immunofluorescence staining, antigen retrieval was performed in a pressure cooker using 0.01 M Tris-EDTA (pH 9.0) buffer. The sections were blocked with 10% goat serum and 1% BSA in TBS for 1 hour at room temperature and incubated with primary antibodies in 1% BSA in TBS overnight at 4°C. The next day, the sections were incubated with Cy2, Cy3 or Cy5-conjugated fluorescent secondary antibodies (Jackson Laboratory, 1:500) and afterwards, DAPI in mounting media was added (Invitrogen #P36930). The following antibodies were used: anti-Ki67 (Abcam #ab15580, 1:200), anti-E-cadherin (BD Biosciences #610182, 1:200), anti-E-cadherin (R&D Systems #AF748) and anti-Cleaved-caspase-3 (Cell Signaling Technology #9661,1:500).

## Western blot analysis

Tumoroids in 3D were released from Matrigel with Cell Recovery Solution (Corning #354253). The tumoroids were then washed with PBS and lysed with RIPA Buffer (Abcam ab156034) supplemented with a protease/phosphatase inhibitor cocktail (Cell Signaling Technology #5872S). Lysate was briefly sonicated and spun down for 15 min at 14000g at 4°C. Supernatant was transferred to a clean tube and protein concentration was determined by BCA assay (Pierce #23225). 4x loading buffer (Biorad #1610747) was then added to the cell lysates and incubated at 98°C for 5 min. Equal amounts of protein were loaded into a 10% SDS–PAGE gel and after transfer, membranes were blocked with 5% milk TBST for 1 hour at room temperature and then incubated with indicated primary antibodies diluted in 5% milk TBST overnight at 4°C. The next day, after washing, membranes were incubated with HRP conjugated secondary antibodies for 1 hour. Signals were detected by HRP-conjugated secondary anti-rabbit antibody (CST#7074S, 1:1,000) and anti-mouse antibody (CST#7076S, 1:2000) visualized with the SuperSignal West Pico PLUS Chemiluminescent Substrate (Thermo Scientific, #34577) and Bio-RAD Chemidoc TMMP imaging system. The following primary antibodies were used: anti-β-ACTIN (Abcam #ab6276, 1:5000), anti-CTNNA1 (Cell Signaling Technology #3236T, 1:1000), anti-BCL2L13 (Proteintech #16612-1AP, 1:1000), and anti-TOM20 (Cell Signaling Technology #42406S, 1:1000).

## Annexin V/PI staining

Tumoroids were digested into single cells and then 2000 cells/well were seeded in triplicate into 24-well plates or low-adherence plates. Detached and attached tumoroids were then cultured for 1 or 2 days respectively and then digested into single cells and stained with Dead Cell Apoptosis Kit with Annexin V Alexa Fluor 488 & PI (Invitrogen #V13241) according to the manufacturer’s instructions. An LSR Fortessa (BD Biosciences) was used to analyze Annexin V and PI staining. The data was then quantified using Flowjo (BD Biosciences).

## EdU incorporation assays

Tumoroids were digested into single cells and then 1000 cells/well were seeded in triplicate into 24-well plates with complete medium supplemented with Y-27632. Tumoroids were cultured for 4 days, after which 10 μM EdU was added to the culture media for 1.5 hours. Tumoroids were digested into single cells and stained with eBioscience Fixable Viability Dye 780 (Invitrogen #65-0865-14). Afterwards, EdU staining was performed with Click-iT Plus EdU Alexa Fluor 647 Flow Cytometry Assay Kit (Thermo Fisher #10634) in accordance with the manufacturer’s instructions. DNA was stained using Fxcycle Violet Stain (Thermo Scientific #F10347). An LSR Fortessa (BD Biosciences) was used to analyze EdU incorporation and DNA content of live cells. The data was subsequently quantified using Flowjo (BD Biosciences).

## Live cell imaging

Black Cellstar 96-well culture plate (Greiner #655087) wells were coated with Matrigel diluted 3:1 in PBS and incubated for 15 min at 37°C. 2000 single tumoroid cells were then seeded on top of Matrigel in complete media and allowed to form tumoroids for 48h. The resulting tumoroids were then imaged on an ImageXpress Micro 4 for 24 hours at 37°C, 20% O_2_. Images were processed into movies and tumoroids tracks were determined by the Fiji plug-in TrackMate^69^. Resulting tumoroid tracks were further analyzed using the celltrackR package.

## scRNA-seq library preparation and analysis

Primary tumors were minced and incubated with 1U/ml Dispase (STEMCELL Technologies #07923) supplemented with 0.05 mg/ml liberase (Sigma-Aldrich# 05401127001) for 1 hour at 37°C. Digested cells were then filtered through 70µM strainers, spun down, and resuspended in Red Blood Cell Lysing Buffer Hybri-Max (Sigma-Aldrich #R7757) for 5 min at RT. Cells were then spun and washed with 0.04%BSA-DPBS and then stained with Cell Multiplexing Oligo (CMO, 10X Genomics #1000261) and incubated for 5 min at room temperature. After washing per the 10x Genomics protocol, cells were resuspended in 0.04% BSA-DPBS supplemented with DAPI and FACS was used to isolate live single cells. The 10x Chromium Next GEM Single Cell 3’ Reagent Kits v3.1 (Dual Index) was used for library preparation according to manufacturer’s instructions (10x Genomics), 2000-5000 cells were partitioned into Gel Beads in Emulsion (GEMs), lysed, and barcoded by reverse transcription of mRNA into cDNA, followed by amplification, enzymatic fragmentation and 5’ adaptor and sample index attachment. Sample libraries were sequenced on the NovaSeq 6000 System (Illumina).

Quantification of raw sequencing data was performed using Cellranger (6.1.2) multi function in conjunction with CMO sequences and refdata-gex-mm10-2020-A reference. Seurat^70^ (4.3.0) was used for standard QC, normalization, and multiple clustering techniques (PCA, TSNE, UMAP) following the appropriate Seurat workflows and their parameters. M1/M2 gensets MM749 and MM750^71^ were imported, and where appropriate converted between organisms using gProfiler^72^ Gene ID conversion web tool, and then used with Seurat’s AddmoduleSore function, to test enrichment of the genesets across our dataset. EnrichR^73^ package’s DEenrichRPlot function was used with MsigDB’s Hallmark 2020 database to test enrichment of hallmark gene signatures across our dataset, with ‘max.genes’ parameter set to 300.

## Macrophage flow cytometry

Primary tumors were first dissected, minced and incubated in 1U/ml dispase (STEMCELL Technologies# 07923) supplemented with 0.05 mg/ml liberase (Sigma-Aldrich# 05401127001) at 37°C for 1 hour. Digested tumors were then washed and filtered through 70µM strainers. Cells were then distributed into a 96 U-Bottom plate and blocked on ice for 10 min with anti-CD16/CD32 solution. A cocktail of antibodies was then added to the cells and incubated for 30 min on ice. The cocktail included anti-Ly6c (Thermo-Fisher 45-5932,1:200), anti-MHC-II (BD Biosciences 562564, 1:200), anti-CD3e (BioLegend 100353, 1:200), anti-CD19 (BD Biosciences 562956, 1:200), anti-CD206 (BioLegend 141723, 1:200), anti-Ly6G (BioLegend 127645, 1:200), anti-CD45.2 (BioLegend 109822, 1:200), anti-Siglec-F (BD Biosciences 552126, 1:200), anti-CD11b (Thermo Fisher 15-0112-82, 1:200), anti-CD64 (Biolegend 139314, 1:200). Live-dead ef780 (eBiosciences 65-0865-18, 1:1500) was also included. Cells were then fixed, permeabilized and washed (BD Biosciences 554714). An LSR Fortessa (BD Biosciences) was used to analyze the stained cell population and the data was then quantified using Flowjo (BD Biosciences).

## RT-PCR and qPCR

For gene expression analysis, tumoroids/monolayers were lysed in TRIzol (Invitrogen# 15596026) and total RNA was isolated according to manufacturer’s instructions. 1.5 µg total RNA was used to synthesize cDNA using the High-Capacity cDNA Reverse Transcription Kit (Applied Biosystems #4368813) following the manufacturer’s instructions. Quantitative PCR reactions (qPCR) using SYBR green (Applied Biosystems #4367659) and QuantiStudio 7 Flex RealTime PCR (Applied Biosystems) was performed. Specificity of primers used was verified by melt curves. Data was normalized by *Gapdh*. The following sets of primers were used to assess *Cdh1*, *Cdh2*, *Snai1*, *Twist1*, *Vim*, *Zeb1*, and *Zeb2* mRNA expression levels: Gapdh, 5′-AGACGGCCGCATCTTCTT-3′ and 5′-TTCACACCGACCTTCACCAT-3′; Cdh1, 5’-CAGTTCCGAGGTCTACACCTT-3’ and 5’-TGAATCGGGAGTCTTCCGAAAA-3’; Cdh2, 5’-AGGCTTCTGGTGAAATTGCAT-3’ and 5’-GTCCACCTTGAAATCTGCTGG-3’; Snai1, 5’-TCCACACGCACCTACAGTCT-3’ and 5’-CCGAGGACCGGGTCACATA-3’; Twist1, 5’-GGACAAGCTGAGCAAGATTCA-3’ and 5’-CGGAGAAGGCGTAGCTGAG Vim, 5’-CACACGCTGCCTTGTGTCT-3’ and 5’-GGTCAGCAAAAGCACGGTT-3’; Zeb1, 5’-ACCGCCGTCATTTATCCTGAG-3’ and 5’-CATCTGGTGTTCCGTTTTCATCA-3’; Zeb2, 5’-AAACGTGGTGAACTATGACAACG-3’ and 5’-CTTGCAGAATCTCGCCACTG-3’;

To determine mitochondrial DNA: nuclear DNA ratios (mtDNA:nDNA), total DNA was isolated from attached or detached tumoroids using QIAamp DNA Micro Kit (Qiagen #56304) according to the manufacturer’s instructions. qPCR was performed on purified DNA using SYBR green (Applied Biosystems #4367659) and QuantiStudio 7 Flex RealTime PCR (Applied Biosystems). Primers targeting 16S (*mt-Rnr2*) and Nd1 (*mt-Nd1*) were used to quantify mtDNA and primers targeting *Hk2* were used to quantify nDNA: 16s, 5’-CCGCAAGGGAAAGATGAAAGAC-3’ and 5’-TCGTTTGGTTTCGGGGTTTC-3’; Nd1, 5’-CTAGCAGAAACAAACCGGGC-3’ and 5’-CCGGCTGCGTATTCTACGTT-3’; Hk2. 5’-GCCAGCCTCTCCTGATTTTAGTGT-3’ and 5’-GGGAACACAAAAGACCTCTTCTGG-3’;

## Tumoroid detachment assays

Tumoroids were digested into single cells and seeded at 2000 cells/well in Nunclon Sphera 96U Well Round Bottom Plates (Thermo Fisher Scientific #174925) using complete medium supplemented with Y-27632, but excluding N-acetyl-cysteine. Detached cells were treated with 20μM Q-VD-OPH (Cayman #15260) as indicated. The CellTiter-Glo viability assay (Promega #G9681) was used according to the manufacturer’s instructions to quantify cell viability 72 hours after seeding. CellTiter-Glo luminescence was read using a Biotek Synergy 2 microplate reader.

## Software and statistical analysis

PRISM software and R were used for statistical analysis, data visualization, and data processing. (http://www.graphpad.com). The R language and environment for graphics (https://www.r-project.org) was used in this study for the bioinformatics analysis of CRISPR screen and scRNA-seq datasets. The R packages used for all analysis described in this manuscript were from the Bioconductor and CRAN, including enhancedVolcano^74^. enrichR^73^ was used for gene set enrichment analysis. Fiji was used for live imaging (Trackmate^69^) and transwell analysis. Seurat^70^ and gProfiler^72^ were utilized for scRNA-seq analysis.

## Supplemental Information

### Supplemental Table Titles

Table S1. Primary screen library sgRNA sequences

Table S2. Transwell screen raw and normalized read counts

Table S3. Transwell screen MAGeCK analysis

Table S4. Primary *in vivo* screen raw and normalized read counts

Table S5. Primary *in vivo* screen processed data organized by mouse

Table S6. Primary *in vivo* screen MAGeCK analyses of primary tumors compared to baseline sample

Table S7. Secondary *in vivo* screen library sgRNA sequences

Table S8. Secondary *in vivo* screen raw and normalized read counts

Table S9. Secondary *in vivo* screen processed data organized by mouse

Table S10. Secondary *in vivo* screen MAGeCK analyses of primary tumors compared to baseline sample

Table S11. Tertiary *in vivo* screen library sgRNA sequences

Table S12. Tertiary *in vivo* screen raw and normalized read counts

Table S13. Tertiary *in vivo* screen processed data organized by mouse

Table S14. *In vivo* screens hit lists

### Supplemental Video Titles

Video S1. Continuous live cell imaging of *sgCtnna1* APK organoids over 24 hours

Video S2. Continuous live cell imaging of *sgBcl2l13* APK organoids over 24 hours

Video S3. Continuous live cell imaging of *sgSafe* APK organoids over 24 hours

**Figure S1.**
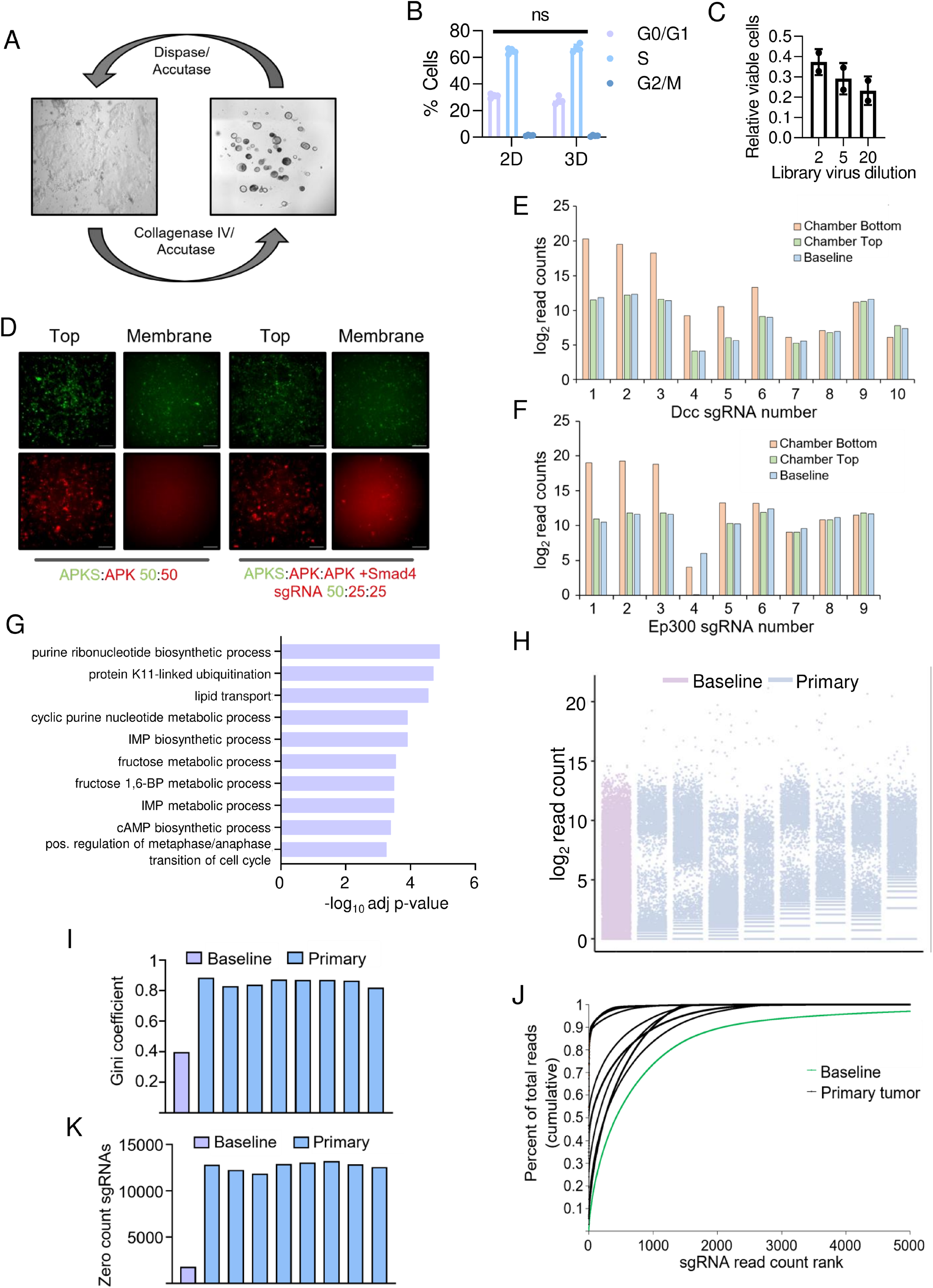
Adaptation of monolayer culture, transwell establishment and primary tumor analysis of main screen. **A.** Brightfield images depicting interconversion between monolayer and organoid/tumoroid cultures. **B.** Quantification of cell cycle status of *APK* monolayer and tumoroid cultures determined by EdU incorporation assays. (n = 3 per group). Statistical significance was assessed via two-way ANOVAs. ns: no significance. **C.** Proportion of viable, sgRNA library-transduced cells after antibiotic selection. Cells were normalized by unselected control. (n = 2 per group). **D.** Representative immunofluorescent images of mixed *APKS-zsGreen* and *APK-mCherry* +/− *Smad4* sgRNA tumoroids in transwell top and membrane respectively. Cells were mixed according to ratios shown. Scale bar: 100µm. **E,F.** Log_2_ normalized read counts of sgRNAs targeting *Dcc* **(E)** and *Ep300* **(F)** in chamber membrane/bottom, top and baseline samples. **G.** GO pathway analysis of a random gene list selected from the sgRNA library. **H.** Log_2_ normalized read counts of library sgRNAs of baseline and primary tumor samples. **I.** Gini coefficients of baseline and primary tumor samples. **J.** Cumulative normalized sgRNA read count distribution of baseline and primary tumor samples. **K.** Number of library sgRNAs with 0 reads in baseline and primary tumor samples. All error bars represent mean ± SD.

**Figure S2.**
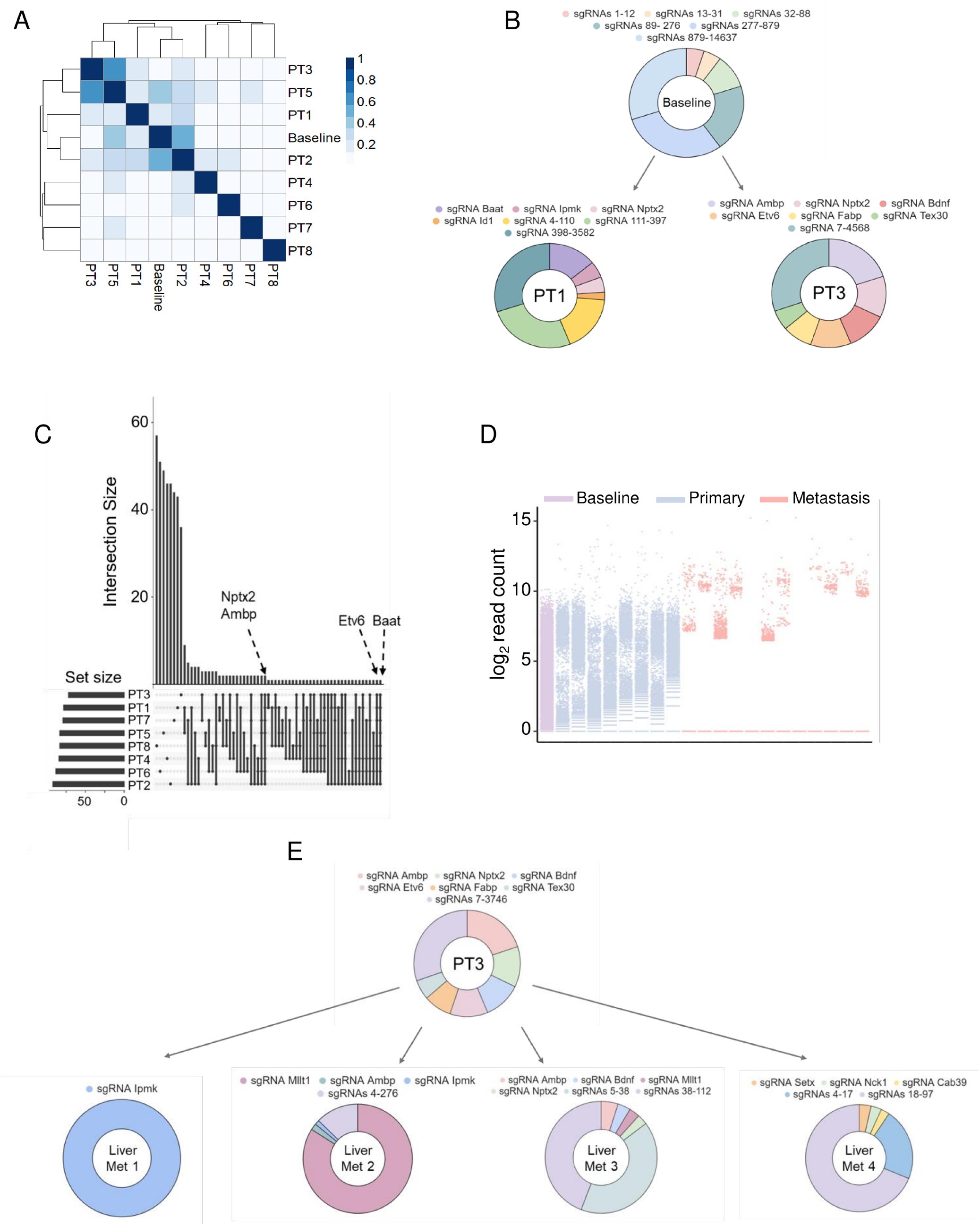
Primary tumor and metastatic analysis of main screen. **A.** Unsupervised clustering of baseline and primary tumor sample sgRNA read count distributions. **B.** Representative sgRNA composition of baseline and primary tumor samples **C.** Overlap of significantly enriched hits called by MAGeCK (*p* < 0.05) in primary tumors relative to baseline sample. *Genes recurring in ≥ 7 primary tumors are highlighted.* **D.** log_2_ normalized read sgRNA counts of baseline, primary and metastatic tumor samples. **E.** Representative sgRNA makeup of primary and metastases isolated from a single mouse.

**Figure S3.**
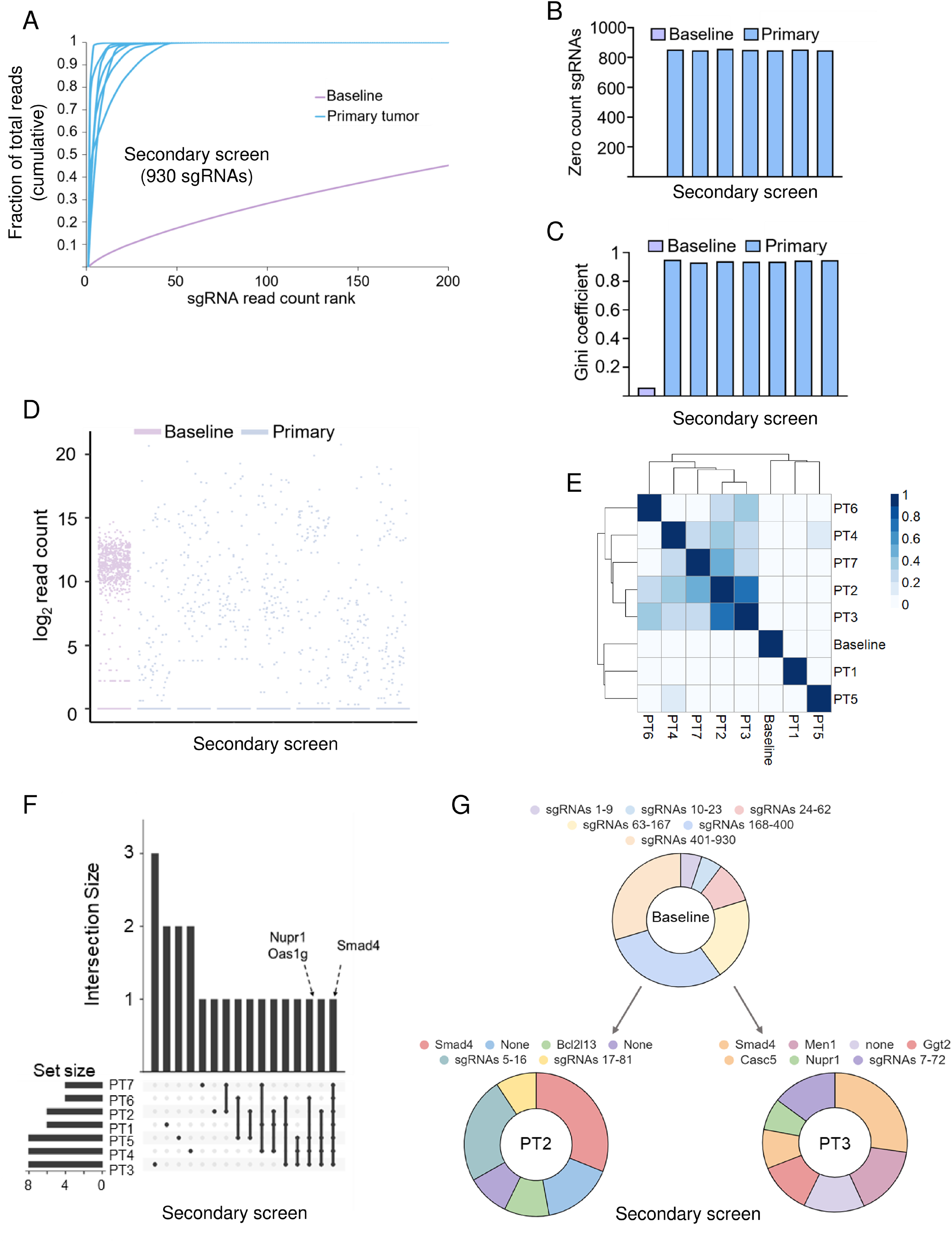
Primary tumor analysis of secondary (930 sgRNA) validation screen. **A.** Cumulative normalized sgRNA read count distribution of baseline and primary tumor samples from the 930 sgRNA secondary screen pool. **B-D.** Number of library sgRNAs with 0 reads **(B),** Gini coefficients **(C)** and log_2_ normalized read counts of baseline and primary tumor from secondary screen sgRNA pool **(D)**. **E.** Unsupervised clustering of baseline and primary tumor sample sgRNA read distributions. **F.** Overlap of significantly enriched hits called by MAGeCK (*p* < 0.05) in primary tumors relative to baseline sample. Genes scoring in ≥ 4 primary tumors are highlighted. **G.** Representative sgRNA composition of baseline and primary tumor samples highlighting enrichment of sgRNAs targeting *Smad4*.

**Figure S4.**
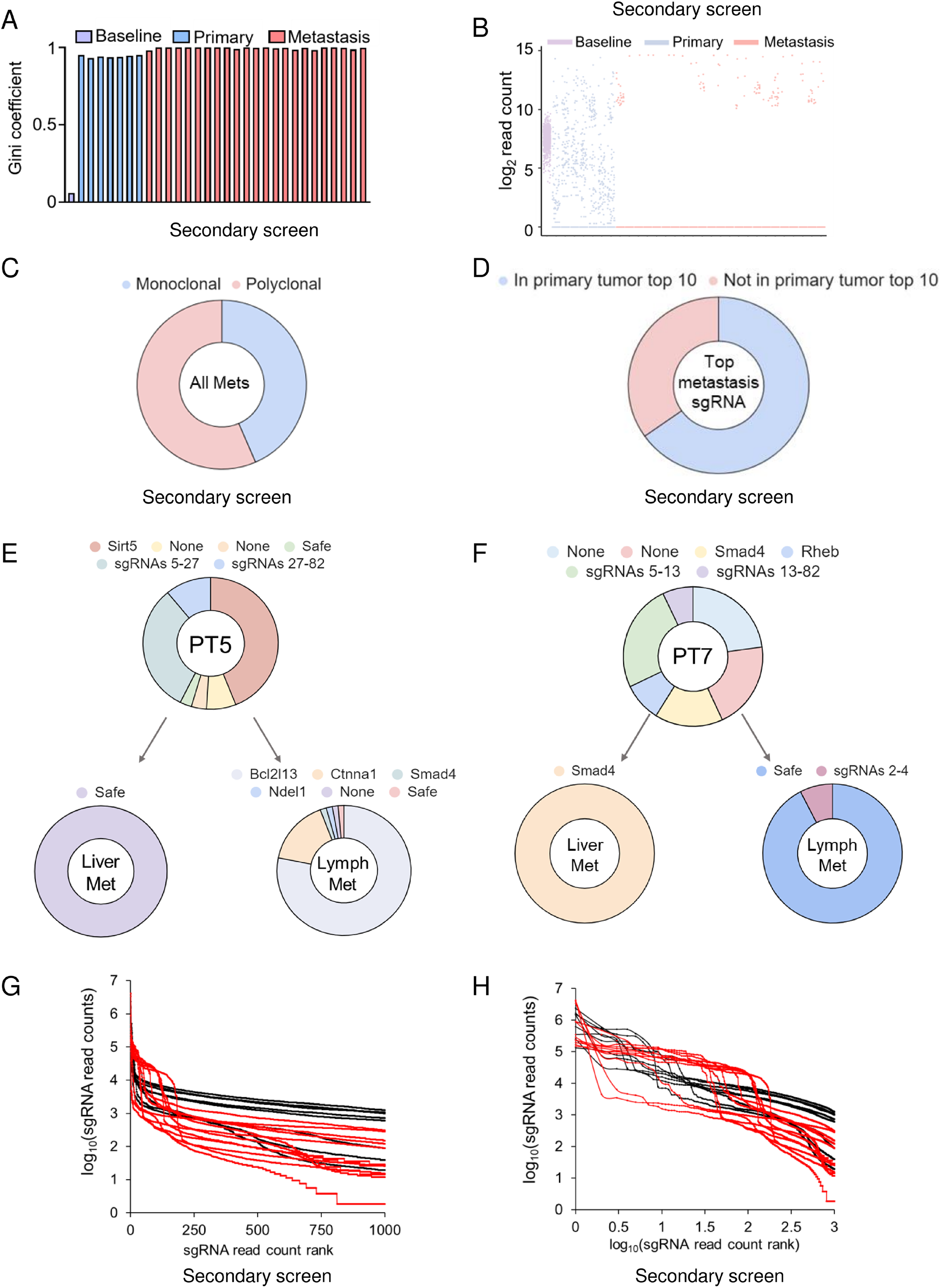
Clonal composition and representative examples of secondary screen metastases. **A,B.** Gini coefficients **(A)** and log_2_ normalized read sgRNA counts **(B)** in baseline, primary tumor, and metastasis samples from secondary (930 sgRNA) screen after filtering. **C.** Clonality of metastases collected in the secondary screen. **D.** Proportion of metastases with the top sgRNA ranking within the top 10 sgRNAs of matching primary tumor. **E, F.** Representative sgRNA composition of matched primary tumors and metastases in the secondary screen highlighting enrichment of candidate suppressor gene *Bcl2l13* **(E)** and positive control *Smad4* **(F)**. **G,H.** sgRNA rank and sgRNA read count data transformations to test possible distributions of read count data. If log transformation of read count data (y-axis) linearizes the relationship between sgRNA read count and sgRNA rank, the read count data fit an exponential distribution (tested in **(G)**). If log-log transformations of both axes linearizes the relationship between sgRNA read count and sgRNA rank, the read count data fit a power law distribution (tested in **(H)**).

**Figure S5.**
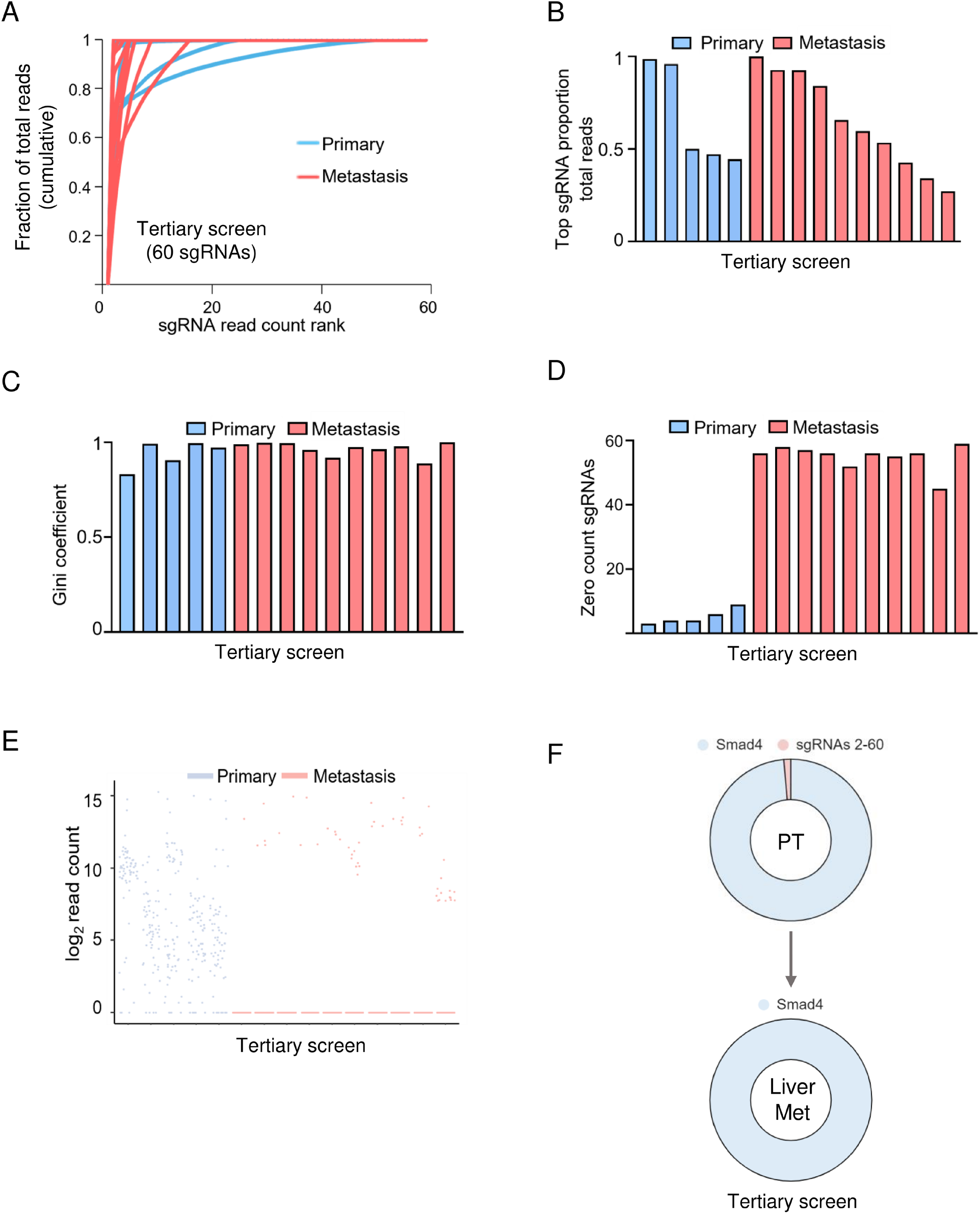
Tertiary (60 sgRNA) screen uncovers drastic bottlenecking accompanying low library diversity. **A.** Cumulative normalized sgRNA read count distribution of tertiary (60 sgRNA) screen baseline, primary tumor and metastasis samples. **B.** The proportion of the total reads mapped to the highest-ranked sgRNA of each primary and metastatic tumor. **C-E.** Gini coefficients **(C)**, number of sgRNAs with 0 reads **(D)** and log_2_ normalized read sgRNA counts **(E)** of tertiary screen baseline, primary tumor and metastasis samples. **F.** Representative sgRNA composition of primary tumor and a matched metastasis highlighting enrichment of sgRNAs targeting *Smad4* in the PT and the metastatic lesion.

**Figure S6.**
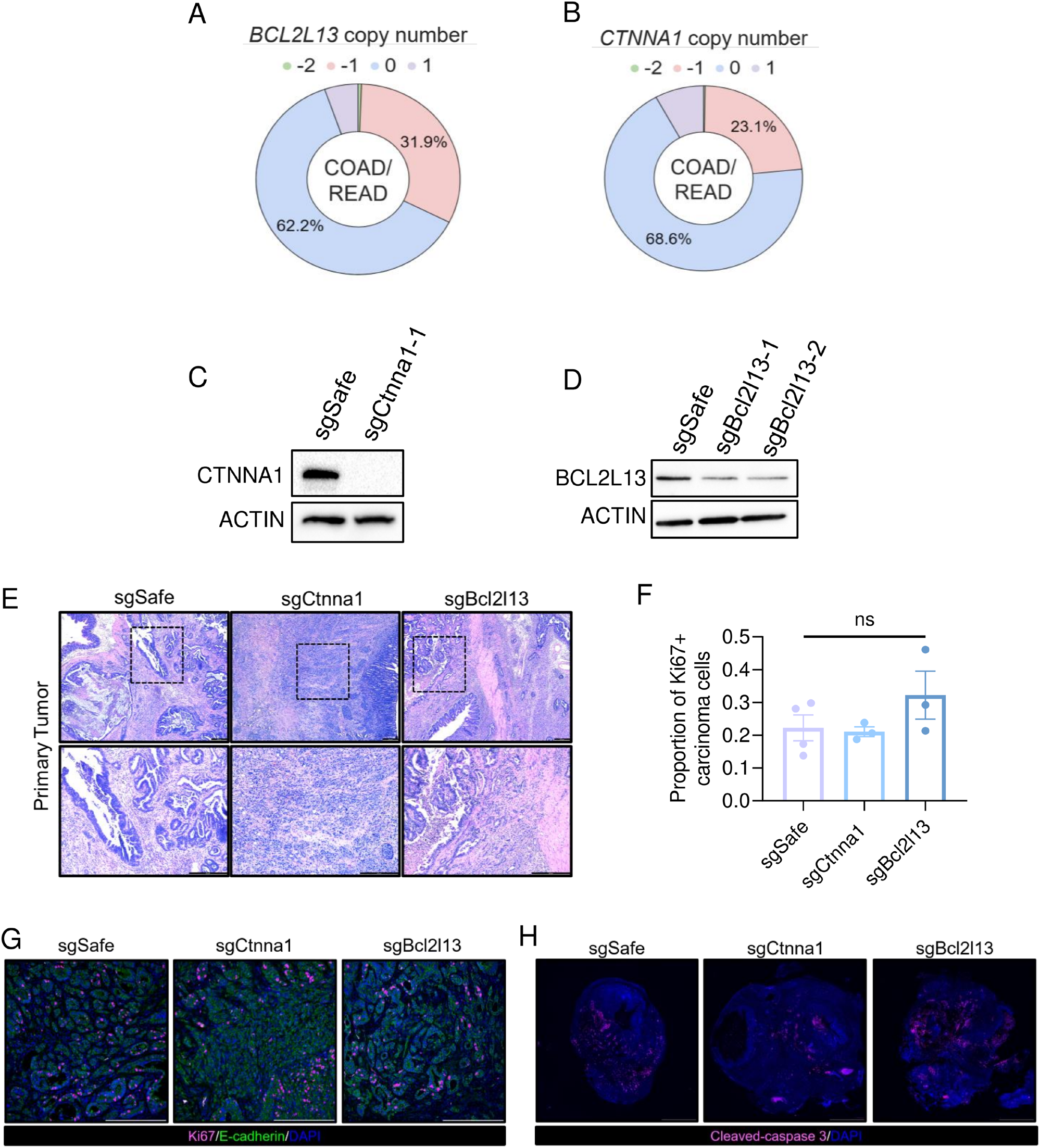
Histological analyses of single-gene validation and copy number analysis. **A, B.** Percentage of COAD/READ patients with the indicated Gistic-thresholded copy number of *BCL2L13* **(A)** and *CTNNA1* **(B) C,D.** Western blot analysis for CTNNA1 **(C)** and BCL2L13 **(D)** in *APK* tumoroids transduced with indicated sgRNAs. β-ACTIN was used as a loading control. **E.** Representative hematoxylin and eosin-stained sections derived from primary tumors formed by orthotopically implanted *APK* organoids transduced with the indicated sgRNAs. Scale bar= 250 µm. **F,G.** Quantification **(F)** and representative images **(G)** of primary tumors stained with antibodies targeting Ki67 (pink) and E-cadherin (green) as well as DAPI (blue) (*n ≥* 3 mice in each group). Scale bar= 250 µm. Error bars represent mean ± SEM. Student’s t-test with Holm-Šídák correction used to assess statistical significance. ns: not significant. **H.** Representative images of primary tumors stained with antibodies targeting cleaved-caspase 3 (pink) as well as DAPI (blue). Scale bar= 2 mm.

**Figure S7.**
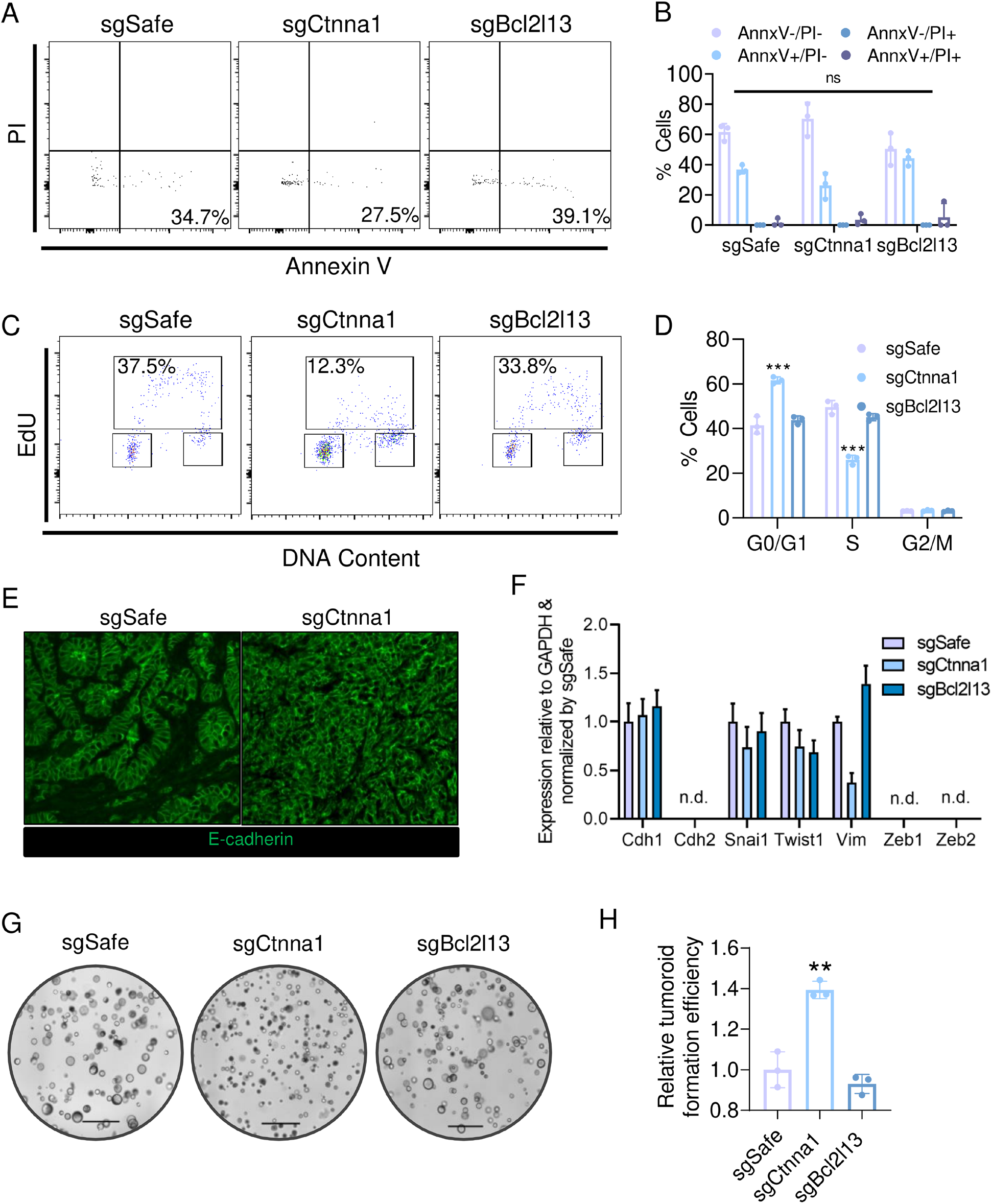
*In vitro* analyses of *sgCtnna1* and *sgBcl2l13 APK* tumoroids. **A, B.** Representative flow cytometry plots **(A)** and corresponding quantification **(B)** of Annexin V and propidium iodide (PI) staining of *APK* tumoroids transduced with *sgSafe* or *sgBcl2l13-1* (n = 3 in each group). Statistical significance assessed via two-way ANOVA with Holm-Šídák multiple comparisons correction. **C,D.** Representative flow cytometry plots **(C)** and quantification **(D)** of EdU incorporation assays in *APK* tumoroids transduced with *sgSafe*, *sgCtnna1-1* or *sgBcl2l13-1*. (n = 3 in each group). Statistical significance assessed via two-way ANOVA with Holm-Šídák multiple comparisons correction. **E.** E-cadherin (green) staining of *APK* primary tumors transduced with the indicated sgRNAs. **F.** RT-qPCR expression profiling of canonical EMT marker genes in *APK* organoids transduced with *sgSafe*, *sgCtnna1-1* or *sgBcl2l13-1*. GAPDH was used as a housekeeping gene and samples were normalized by *sgSafe* sample (*n* = 3 each group). n.d.: not detected. Error bars represent mean ± SEM. **G,H.** Representative brightfield images **(G)** and quantification **(H)** of tumoroid formation assays in *APK* tumoroids transduced with the indicated sgRNAs. (n = 3 in each group). Statistical significance assessed via student’s t-test with Holm-Šídák correction. For all panels except F, error bars represent mean ± SD. For all panels, ** *p* < 0.01, *** *p <* 0.001.

**Figure S8.**
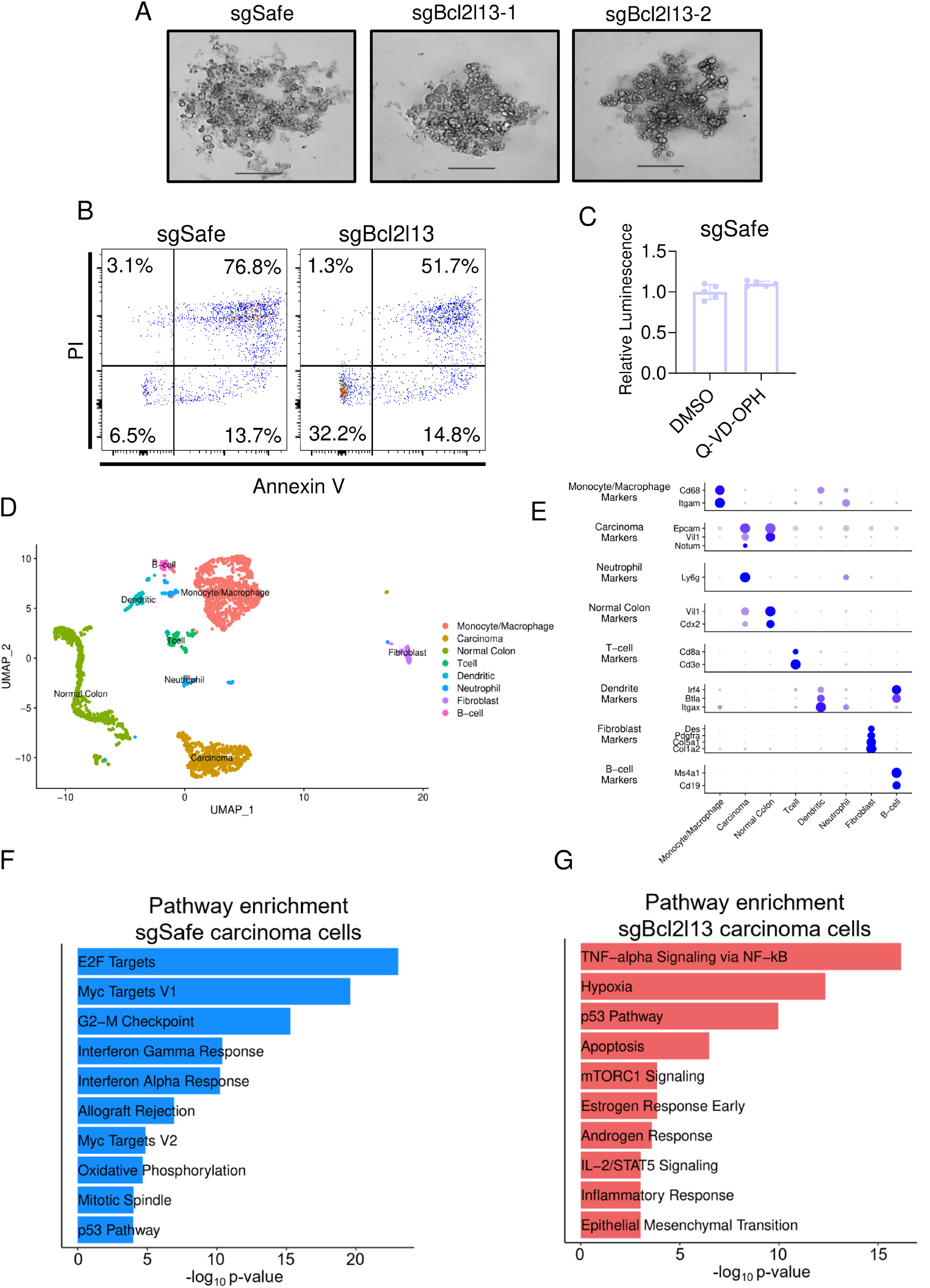
BCL2L13 regulates *APK* tumoroid survival after extracellular matrix detachment and scRNA-seq analysis of sgBcl2l13 primary tumors. **A.** Representative brightfield images of *APK* tumoroids transduced with the indicated sgRNAs after 72h detachment. Scale bar = 200µm. **B.** Representative flow cytometry plots of Annexin V and propidium iodide (PI) staining of *sgSafe* or *sgBcl2l13-1 APK* tumoroids transduced with the indicated sgRNAs after 24h detachment. **C.** CellTiter-Glo assay luminescence normalized readings of *sgSafe APK* tumoroids treated with 20μM Q-VD-OPH for 72h. Error bars represent mean ± SD. **D.** Uniform Manifold Approximation and Projection (UMAP) of all cells captured in scRNA-seq of *sgSafe* and *sgBcl2l13* primary tumors. **E.** Marker genes used for cell annotation in **(D)**. **F.** Pathways downregulated in *sgBcl2l13* carcinoma cells according to GSEA. **G.** Pathways upregulated in *sgBcl2l13* carcinoma cells according to GSEA.

**Figure S9.**
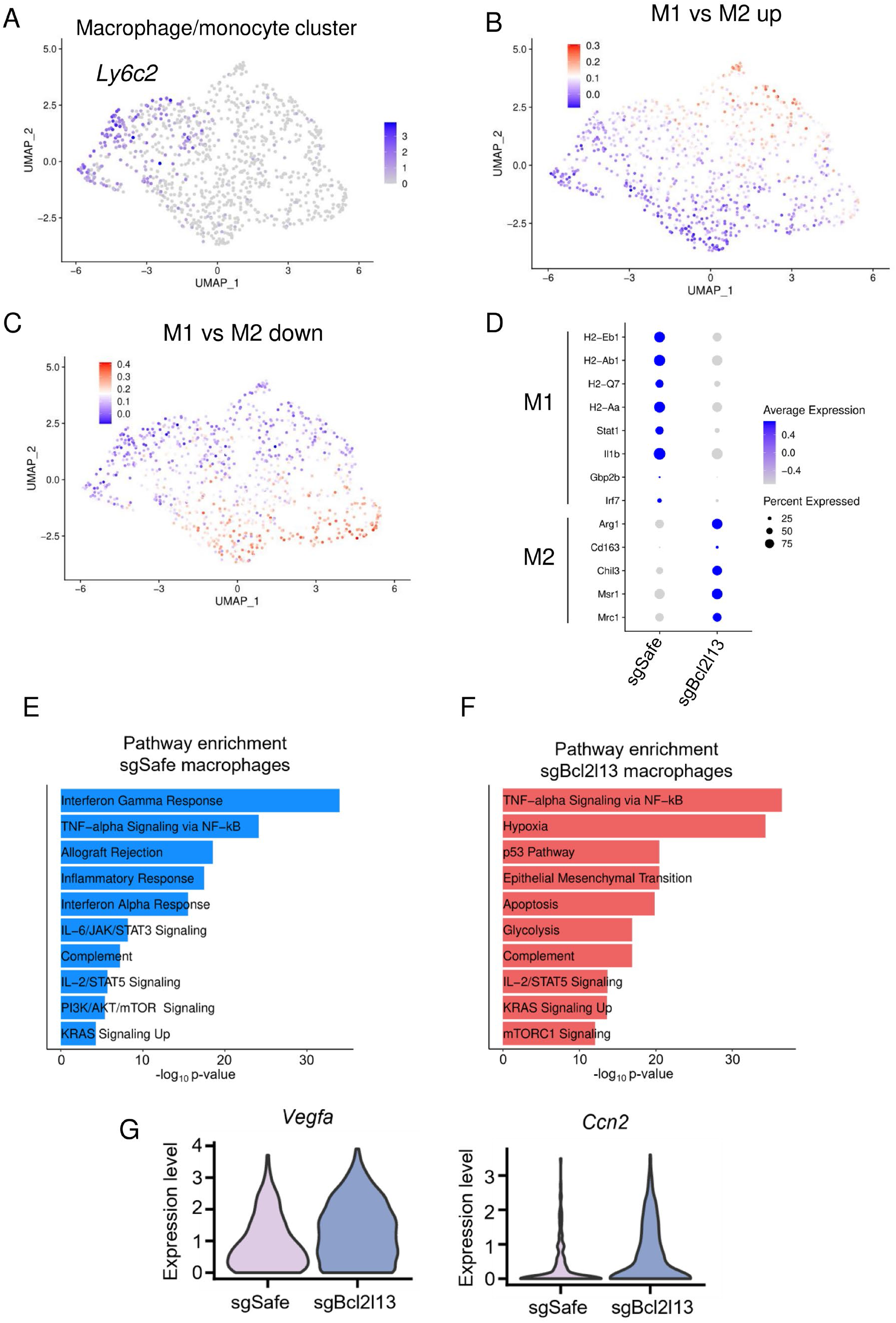
M1 and M2 signatures in *sgSafe* and *sgBcl2l13* macrophages. **A.** *Ly6c2* expression in macrophage/monocyte cluster from scRNA-seq of *sgSafe* and *sgBcl2l13 APK* primary tumors. **B,C.** Relative Gene set Expression Score of M1-like genes **(B)** and M2-like genes **(C)** on Uniform Manifold Approximation and Projection (UMAP) of macrophages/monocytes calculated using Seurat. **D.** Expression of individual M1-like and M2-like genes in *sgSafe* and *sgBcl2l13* primary tumor monocytes/macrophages. **E.** Pathways downregulated in *sgBcl2l13* macrophages according to GSEA. **F.** Pathways upregulated in *sgBcl2l13* macrophages according to GSEA. **G.** Expression of *Ccn2* and *Vegfa* in *APK* PT carcinoma cells.

**Figure S10.**
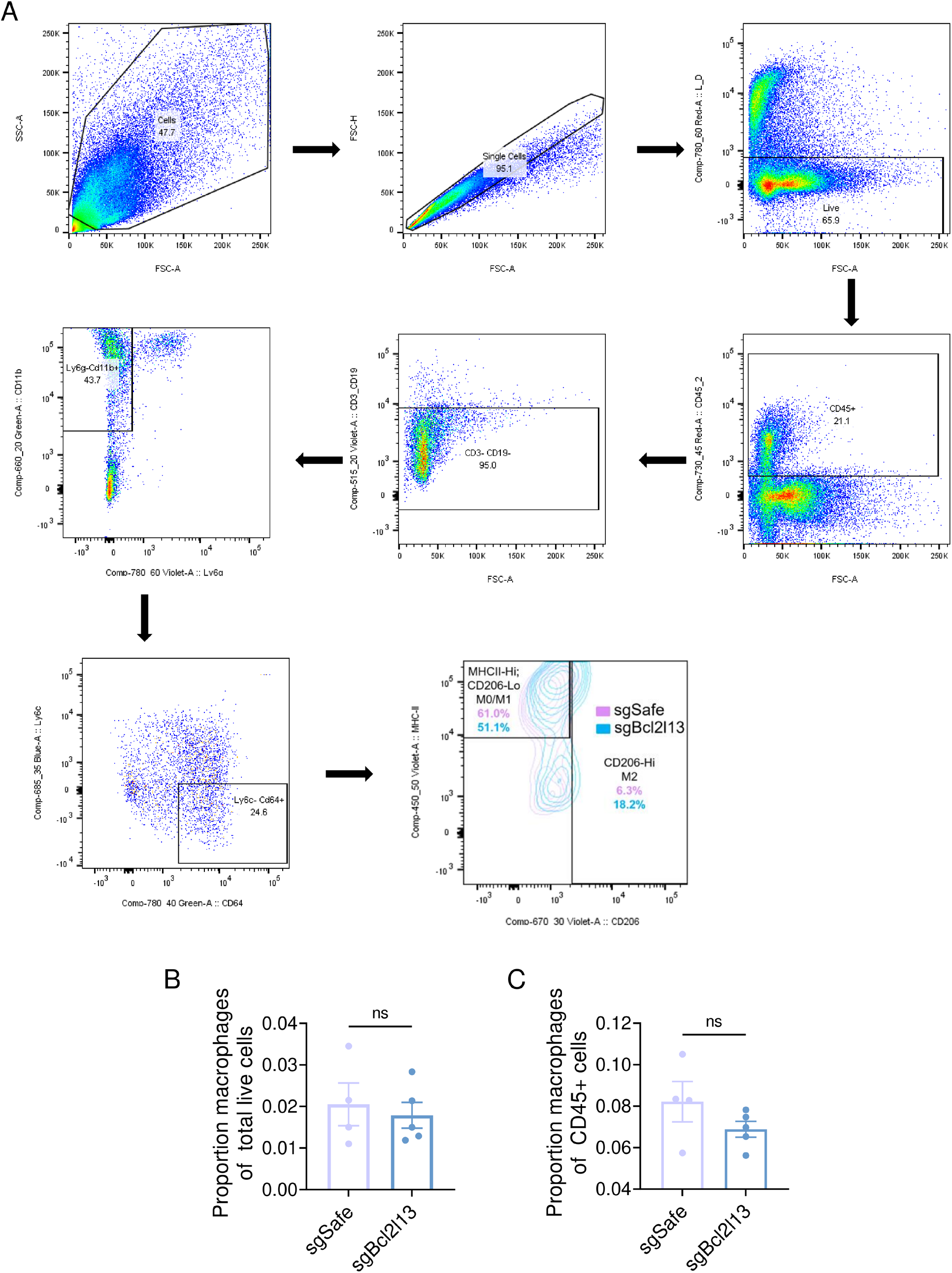
Gating scheme and quantification of flow cytometric analysis of *APK* primary tumors. **A.** Macrophages were defined as viable CD45+;CD3-;CD19-;LY6G-;CD11b+;LY6C-;CD64+ cells that were MHC-II-Hi and/or CD206-Hi. **B, C.** Proportion of macrophages in *sgSafe* and *sgBcl2l13 APK* primary tumors relative to total live cells **(B)** or total CD45+ cells **(C)** (*n ≥* 4). For all panels, Student’s t-test. Error bars represent mean ± SEM.

## Notes

### Competing Interest Statement

The authors have declared no competing interest.

